# Identification of NLR-associated amyloid signaling motifs in filamentous bacteria

**DOI:** 10.1101/2020.01.06.895854

**Authors:** Witold Dyrka, Virginie Coustou, Asen Daskalov, Alons Lends, Thierry Bardin, Mélanie Berbon, Brice Kauffmann, Corinne Blancard, Bénédicte Salin, Antoine Loquet, Sven J. Saupe

## Abstract

NLRs (Nod-like receptors) are intracellular receptors regulating immunity, symbiosis, non-self recognition and programmed cell death in animals, plants and fungi. Several fungal NLRs employ amyloid signaling motifs to activate downstream cell-death inducing proteins. Herein, we identify in Archaea and Bacteria, short sequence motifs that occur in the same genomic context as fungal amyloid signaling motifs. We identify 10 families of bacterial amyloid signaling sequences (we term BASS), one of which (BASS3) is related to mammalian RHIM and fungal PP amyloid motifs. We find that BASS motifs occur specifically in bacteria forming multicellular structures (mainly in *Actinobacteria* and *Cyanobacteria*). We analyze experimentally a subset of these motifs and find that they behave as prion forming domains when expressed in a fungal model. All tested bacterial motifs also formed fibrils *in vitro.* We analyze by solid-state NMR and X-ray diffraction, the amyloid state of a protein from *Streptomyces coelicolor* bearing the most common BASS1 motif and find that it forms highly ordered non-polymorphic amyloid fibrils. This work expands the paradigm of amyloid signaling to prokaryotes and underlies its relation to multicellularity.

## Introduction

NLRs are intracellular receptors controlling innate immunity and host-symbiont interactions, both in plants and animals (Jones et al. 2016; Mermigka et al. 2019). NLR proteins have a typical tripartite architecture with an N-terminal effector domain, a central (NACHT or NB-ARC) nucleotide binding and oligomerization domain and a C-terminal leucine-rich repeat (LRR) domain. Filamentous fungi also display large and diverse repertoires of up to several hundreds NLR-related genes per genome (Dyrka et al. 2014). These fungal NLR homologs however display WD40, ANK or TPR repeats as ligand recognition domains instead of LRRs found in most plant and animals NLRs (Dyrka et al. 2014; Urbach and Ausubel 2017). Fungal NLRs were found to control programmed cell death associated with non-self recognition in several fungal species (Chevanne et al. 2009; Choi et al. 2012; Daskalov et al. 2015b; Espagne et al. 2002; Heller et al. 2018; Saupe et al. 1995). These proteins are considered the fungal counterparts of plant and animal NLRs (Dyrka et al. 2014, Paoletti and Saupe 2009, Uehling et al. 2017). Remarkably some fungal NLRs employ an amyloid signaling mechanism to engage cell death inducing effector proteins (Loquet and Saupe 2017). These NLRs display a short 20-25 amino acid long N-terminal amyloid forming motifs upstream of the NACHT (or NB-ARC) domain while their cognate effector protein displays a similar motif C-terminally. The amyloid fold of the activated NLR receptor serves as a structural template to convert the homologous region in the effector protein to a similar amyloid fold (Cai et al. 2014; Daskalov et al. 2015b; Daskalov et al. 2012). This signaling mechanism based on the prion principle (self-propagation of protein polymers) is also operating in several immune signaling cascades in mammals, albeit with a different underlying structural basis (Cai et al. 2016). In fungi, NLR and their cognate regulated effector proteins are typically encoded by adjacent genes and thus form functional gene pairs or clusters (Daskalov et al. 2015a; Daskalov et al. 2016; Daskalov et al. 2012). Several classes of fungal amyloid signaling motifs have been described. HRAM motifs (for <u>H</u>ET-s <u>R</u>elated <u>A</u>myloid <u>M</u>otifs) (Pfam PF11558) were originally identified in the [Het-s] prion of *Podospora anserina* and have been subject to in-depth functional and structural characterization (Daskalov et al. 2015a; Daskalov et al. 2015b; Wasmer et al. 2008; Wasmer et al. 2010). HRAMs form a β-solenoid amyloid fold comprising 21 amino acid long pseudo-repeats, with one repeat copy on the NLR and two copies on the effector protein. Another family of signaling amyloids is defined by the sigma motif, involved in the propagation of the *σ* cytoplasmic infectious element of *Nectria haematococca* (Daskalov et al. 2012; Graziani et al. 2004) (Pfam PF17046). Finally, the PP-motif (for pseudo-palindrome) was described in *Chaetomium globosum* and displays a sequence similarity with the mammalian RHIM amyloid sequence scaffolding the RIP1K/RIP3K necrosome (Daskalov et al. 2016; Li et al. 2012; Sun et al. 2002). RHIM and RHIM-related motifs were also identified in viruses and Drosophila (Kleino et al. 2017; Pham et al. 2019). The similarity between the metazoan RHIM and fungal PP-motifs raised the possibility of an ancient evolutionary origin of this mechanism of cell death-inducing amyloid signaling (Daskalov et al. 2016; Kajava et al. 2014).

Several types of downstream effector protein domains activated by NLR-mediated amyloid signaling were described (Daskalov et al. 2012; Dyrka et al. 2014) but the most common are part of a family of membrane targeting proteins including the HeLo, HeLL (Helo-like) and SesA domains (Pfam PF14479, PF17111 and PF17707 respectively). The HeLo domain forms a α-helical bundle with an N-terminal hydrophobic helix and functions as a membrane targeting pore-forming domain (Greenwald et al. 2010; Seuring et al. 2012). The fungal HeLo/Helo-like/SesA domain family shows homology with the N-terminal helical cell death execution domain of the MLKL protein controlling mammalian necroptosis and the RPW8 and Rx-N CC-domains regulating plant immune cell death (Daskalov et al. 2016; Murphy et al. 2013). The homology between the membrane-targeting domains suggests a common evolutionary origin for these defense-related programmed cell death processes in plants, animals and fungi. The plant Rx_N and the HET-S HeLo domain share a common mechanism of membrane targeting based on the membrane insertion of a hydrophobic N-terminal α-helix (Seuring et al. 2012; Wang et al. 2019). Other amyloid-controlled effector domains in fungi are predicted to carry enzymatic activity, in particular the SesB α/β hydrolase domain and the PNP-UDP phosphorylase domain (Daskalov et al. 2016; Daskalov et al. 2012). Noteworthy is the observation that fungal NLRs are found in two distinct domain architectures, either as two-component gene clusters involving amyloid signaling or more frequently in an “all-in-one” architecture with the effector/NOD/repeat domains encoded as single polypeptide (Daskalov et al. 2012; Dyrka et al. 2014).

Several bacterial proteins have received a RHIM amyloid motif annotation in the Pfam or InterPro databases (Rebsamen et al. 2009; Sun et al. 2002). We have analyzed the corresponding protein sequences and found that the region annotated as RHIM occurs in a similar genomic context as fungal amyloid signaling motifs. The motif occurs at the N-terminus of a NLR-like protein and at the C-terminus of a putative effector protein encoded by an adjacent gene. Here, we systematically explore in a genome mining approach the occurrence of putative amyloid signaling sequences in bacterial and archaeal genomes and identify ten families of bacterial amyloid signaling sequences (named here BASS1 to 10). The family designated as BASS3 corresponds to the RHIM-like sequence. We find these motifs specifically in multicellular bacteria in particular in *Actinobacteria* and *Cyanobacteria*. We show for motifs of the BASS1 and BASS3 family, prion formation in the *Podospora anserina* fungal model and fibril formation *in vitro*. We use solid-state NMR and X-ray diffraction to structurally characterize the BASS1 motif from *Streptomyces coelicolor* and find that it assembles into highly ordered non-polymorphic amyloids as previously described for fungal amyloid signaling motifs. We propose that NLR-associated BASS motifs are analogous to the amyloid prion motifs identified in fungi and that this signaling mechanism is shared by filamentous fungi and filamentous bacteria.

## Results

### RHIM-like motifs in Bacteria

In the Pfam database, the majority of the sequences with a RHIM annotation are from Metazoan or Metazoan viruses (El-Gebali et al. 2018; Kajava et al. 2014; Kleino et al. 2017; Li et al. 2012; Pham et al. 2019). A few hits (24/286) however occur in Bacteria. We examined these bacterial RHIM-annotated proteins and found that ten of them are homologous proteins with the RHIM annotation occurring C-terminally, downstream of ∼100 amino acid long predicted α-helical domain that we termed Bell (for <u>b</u>acterial domain analogous to H<u>ell</u>, based on shared features with the fungal HeLo-related domains, see below). In the actinobacterium strain *Saccharothrix* sp. ALI-22-I, the gene adjacent to the gene encoding the Bell/RHIM protein (ONI86675.1) encodes a protein with NLR architecture (NB-ARC domain and TPR repeats) (ONI86674.1). Remarkably, the N-terminal region of this NB-ARC/TPR protein shows sequence homology to the C-terminal RHIM-annotated region of the Bell-domain protein (Fig. 1A and B). The ONI86675.1/ ONI86674.1 bacterial gene pair from *Saccharothrix* therefore displays the same features as effector/NLR gene pairs described in filamentous fungi (Daskalov et al. 2015b; Daskalov et al. 2012). This similarity prompted us to analyze further Bell domain proteins in bacterial genomes.

**Figure 1.**
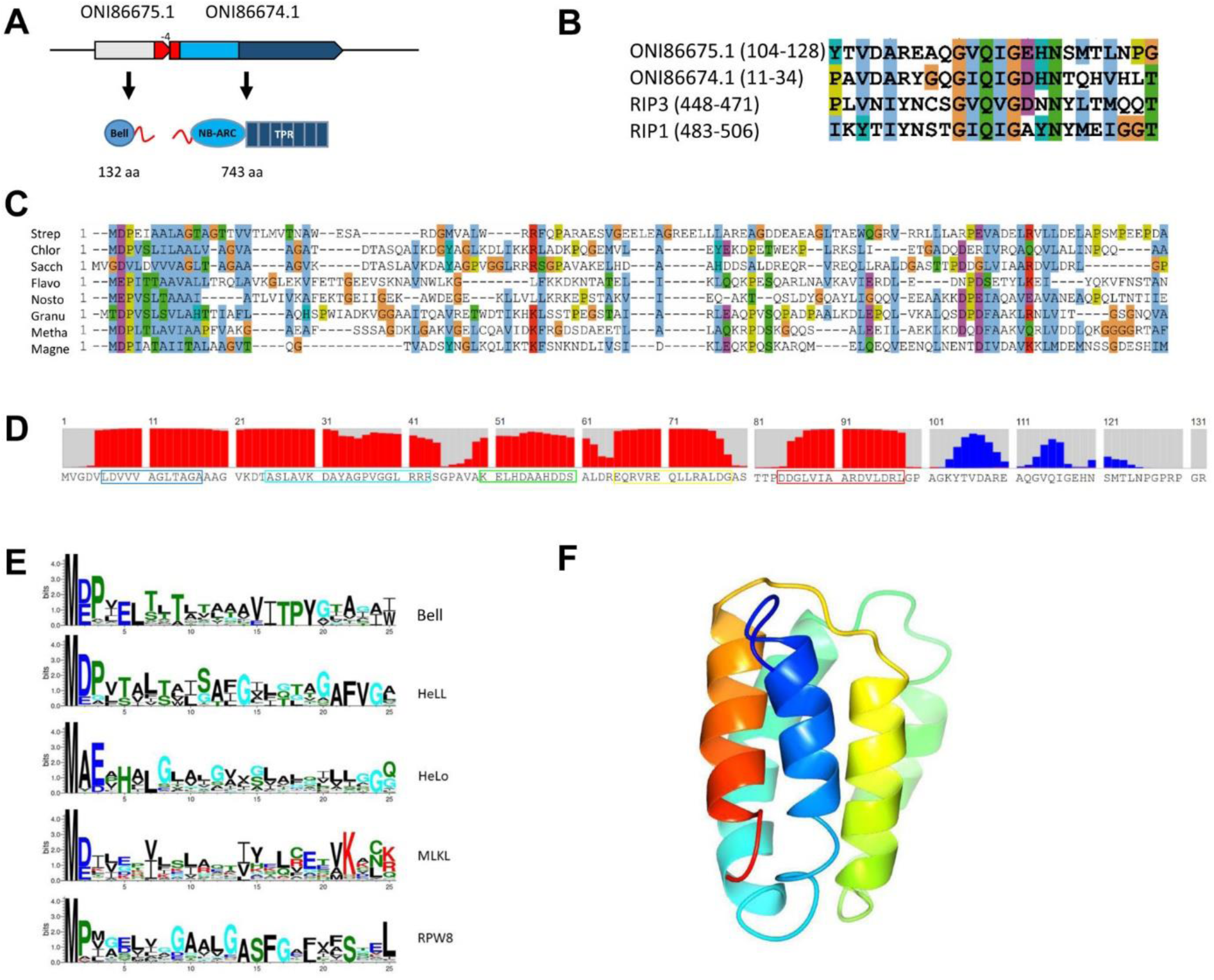
RHIM-like motifs in bacteria are associated to Bell domains. **A.** Genome and domain architecture of the ONI86675.1 and ONI86674.1 gene pair from the actinobacterium strain *Saccharothrix* sp. ALI-22-I. The relative orientation and the overlap between the two ORF are given (the two ORF overlap by 4 bp) as well as the size and domain architecture of the corresponding proteins, respectively for a Bell-domain protein with a C-terminal RHIM-like motif and a NLR-related protein with a NB-ARC and TPR repeat domain and an N-terminal RHIM-like motif. The RHIM-like motif are represented in red in the ORF diagram and the protein cartoon. B. Alignment of the RHIM-like motifs of the proteins encoded by the ONI86675.1 and ONI86674.1 gene pair and the RHIM-motif of the human RIP1 and RIP3 kinases. **C.** Alignment of Bell-domains from various prokaryotes (Strep, Q9RDG0 from Streptomyces coelicolor A3(2); Chlor, HBY96210.1 from *Chloroflexi bacterium*; Sacch, ONI86675.1 from *Saccharothrix sp. ALI-22-I*; Flavo, SDZ50707.1 from *Flavobacterium aquidurense*; Nosto, RCJ33357.1 from *Nostoc punctiforme NIES-2108*; Granu, ROP69996.1 from *Granulicella sp. GAS466*; Metha, AEB69174.1 from *Methanothrix soehngenii (strain ATCC 5969)*; Magne, ETR68090.1 from *Candidatus Magnetoglobus multicellularis str. Araruama*). **D.** Secondary structure prediction for ONI86675.1 from *Saccharothrix sp. ALI-22-I,* red bars represent α-helical propensity, blue bar β-sheet propensity. Boxing corresponds to the α-helices predicted in the homology model given in G. **E.** Consensus sequence of the 25 N-terminal residues of the Bell-domain and other predicted or known cell death execution domains in fungi, plants and mammals. The consensus sequence was generated with Weblogo from a HHMER alignment using the following sequences as queries : ONI86675.1 (Bell), *C. globosum* HELLP (Hell), *P. anserina*, HET-S (HeLo), mouse MLKL (MLKL) and *Arabidopsis thaliana* RPW8.1 (RPW8). **F.** Homology model of the Bell-domain of ONI86675.1 from *Saccharothrix sp. ALI-22-I* based on a contact map (model generated by RAPTOR-X contact).

### Bell-domain occurs in multicellular Bacteria and Archaea

The predicted globular Bell domain of the ONI86675.1 protein from *Saccharothrix* was used as query in HMMER searches to recover homologous proteins. We found that the Bell-domain occurs mainly in short ∼120-140 amino acid long proteins and in more rare instances as N-terminal domain of proteins with a NLR domain architecture (as previously described for HeLo, Helo-like and SesA domains) (Dyrka et al. 2014). Figure 1C gives an alignment of Bell domain proteins from phylogenetically diverse prokaryotes including a sequence from an Archaea (*Methanothrix soehngenii*). The N-terminal region of the domain was predicted to correspond to an hydrophobic α-helix and a HMM-signature of the domain shows frequent occurrence of a negatively charged residue in position 2 or 3, a feature that is common to fungal HeLo, HeLL domains, mammalian MLKL and plant RPW8 and Rx_N domains (Fig.1D, E and F) (Adachi et al. 2019; Daskalov et al. 2016). Using contact prediction maps based on evolutionary co-variance, the Bell domain of *Saccharothrix* sp. ALI-22-I ONI86675.1 was modelled as a five-helix bundle (Fig.1 F). Secondary structure and fold prediction for the different proteins presented in the alignment in Fig. 1C resulted in five-helix bundles for all proteins although in some cases with different topologies (Fig. S1).

We analyzed the phylogenetic distribution of the Bell domain in prokaryotic genomes using the genome-based phylogeny developed by Parks et al. which has been shown to be more accurate than the NCBI taxonomy (Parks et al. 2018). The domain shows a heterogeneous phylogenetic distribution (Table 1, Table2, Table S1). It is most frequent in the *Cyanobacteriota, Actinobacteriota* and *Chloroflexota* but very rare or absent in other bacterial phyla. Within these phyla, the domain is specifically present in genera containing multicellular species (described in Bergey’s Manual of Systematic Bacteriology) (Table S1). In the *Actinobacteria* class, the domain is frequent in families encompassing filamentous species (*Streptomycetaceae, Micronosporaceae, Pseudonocardiaceae, Streptosporangiaceae, Frankiaceae* and *Corynebacteriaceae*) but absent in unicellulars such as the *Bifidobacteriaceae* and *Propionibacteriaceae* (Table 2, Table S1). The same is true at higher phylogenetic resolution, within the *Corynebacteriaceae* family for instance, the domain is highly represented only in the *Nocardia* and *Dietzia* genera encompassing filamentous species (Table S1). In the *Cyanobacteriota*, the domain is highly represented in the *Nostoceceae* family and present in multicellular genera like *Nostoc, Calothrix, Microcystis* or *Fischerella* but absent in the *Cyanobiaceae* family encompassing unicellular genera like *Prochlorococcus* and *Synechococcus* (Table 2, Table S1) (Shih et al. 2013). In the *Chloroflexota*, the domain is specific to the *Chloroflexus* genus encompassing filamentous species like *Chloroflexus auranticus* and *Chloroflexus aggregans*. Similarly, in Archaea, the Bell domain is found exclusively in the *Methanosarcinales* order, in particular in the genus *Methanosarcina* comprising multicellular species forming cell aggregates like *Methanosarcina acetivorans* (Table S1).

**Table 1.** Phylogenetic distribution of <u>Bell domain</u>s and NLRs in Bacteria.

| phylum | #genomes | NBARC-TPR | NACHT-WD | Bell |
| --- | --- | --- | --- | --- |
| d__Bacteria;p__Acidobacteriota | 91 | 9 | 0 | 2 |
| d__Bacteria;p__Actinobacteriota | 13236 | 2533 | 284 | 930 |
| d__Bacteria;p__Bacteroidota | 2858 | 38 | 8 | 30 |
| d__Bacteria;p__Campylobacterota | 3368 | 3 | 0 | 0 |
| d__Bacteria;p__Chloroflexota | 273 | 48 | 9 | 12 |
| d__Bacteria;p__Cyanobacteriota | 513 | 223 | 152 | 173 |
| d__Bacteria;p__Deinococcota | 81 | 6 | 0 | 0 |
| d__Bacteria;p__Desulfobacterota | 267 | 6 | 1 | 3 |
| d__Bacteria;p__Firmicutes | 32747 | 386 | 2 | 0 |
| d__Bacteria;p__Firmicutes_A | 3076 | 34 | 0 | 0 |
| d__Bacteria;p__Firmicutes_B | 130 | 3 | 0 | 0 |
| d__Bacteria;p__Firmicutes_C | 182 | 1 | 0 | 0 |
| d__Bacteria;p__Fusobacteriota | 195 | 0 | 0 | 0 |
| d__Bacteria;p__Myxococcota | 91 | 20 | 18 | 0 |
| d__Bacteria;p__Omnitrophota | 88 | 0 | 0 | 0 |
| d__Bacteria;p__Patescibacteria | 1772 | 10 | 0 | 0 |
| d__Bacteria;p__Planctomycetota | 178 | 4 | 0 | 0 |
| d__Bacteria;p__Proteobacteria | 51737 | 146 | 26 | 83 |
| d__Bacteria;p__Spirochaetota | 808 | 4 | 0 | 0 |
| d__Bacteria;p__Verrucomicrobiota | 582 | 8 | 0 | 0 |

**Table 2.**
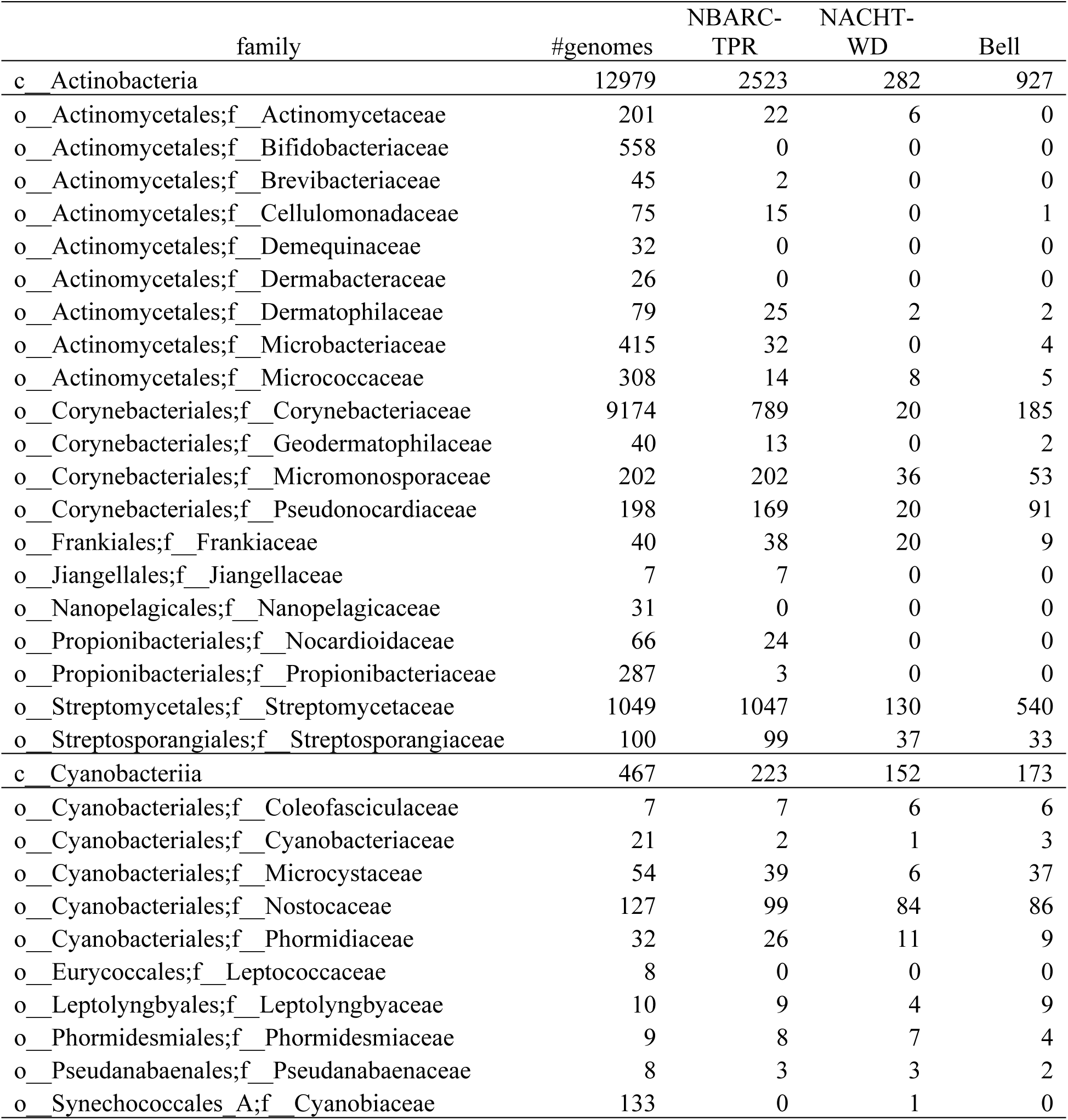
Phylogenetic distribution of Bell-domains and NLRs in Actinobacteria and Cyanobacteria.

The phylogenetic distribution of the Bell domain is mirrored by the distribution of NLR-like architecture proteins (here a NB-ARC or NACHT domain associated with TPR or WD repeats) (Table 1, Table S1). The NLR-like domain architecture is frequent in phylogenetic groups in which the Bell domain is represented (for instance in *Actinobacteriota* and *Cyanobacteriota*) but rare or absent in groups lacking Bell domains (for instance *Proteobacteria* or *Firmicutes*) (Table 1). The same is true at higher phylogenetic resolution (Table 2, Table S1). For instance, in *Cyanobacteriota*, the NLR-like architecture is not found in *Cyanobiaceae* but is prevalent in *Nostocaceae*. Similarly, in Archaea, in *Methanosarcina*, the NLR-like architecture is abundant but rare or absent in other genera. The phylogenetic distributions of NLR-like architectures and Bell domain do not fully overlap though, as for instance the Bell domain is not found in the phylum *Myxococcota* where NLR-like architectures are present. Conversely, the genus *Janthinobacterium* contains many Bell domains but lacks NLR-like architecture proteins (Table S1). Specific occurrence in *Cyanobacteria* and *Actinobacteria* of NLR-type architectures (NB-ARC/NACHT with TPR/WD repeats) was reported previously (Asplund-Samuelsson et al. 2012; Koonin and Aravind 2002; Urbach and Ausubel 2017).

Overall, the Bell domain was found in roughly 1% of the analyzed bacterial genomes (1237 of 113 324) and 10% of the archaeal genomes (94/1183). The domain appears to be typical of Bacteria and Archaea with a multicellular organization either in filamentous or aggregate forming species. In addition, the phylogenetic distribution of the Bell domain proteins is largely mirrored by the phylogenetic distribution of proteins with NLR-like architectures.

### Prediction of amyloid signaling sequence motifs in bacterial genomes

Amyloid signaling motifs in fungi occur in adjacent gene pairs (Daskalov et al. 2012). Based on the observed Bell-RHIM/NLR-like gene pairs (Fig. 1A), we hypothesized that an amyloid signaling mechanism akin to the one described in filamentous fungi might also operate in some bacterial species. To identify potential amyloid signaling motifs in bacteria we extended a genome mining strategy that we have previously developed to identify amyloid signaling motifs in fungi (Daskalov et al. 2012, 2016). We screened for sequence motifs common to C-termini of Bell-domain proteins and N-termini of NLR-like proteins, which are encoded by adjacent (or closely linked) genes. Specifically, we generated a data set of 1816 non-redundant C-termini of prokaryotic Bell-domain proteins, searched for conserved gapless motifs using MEME and identified 117 motifs. For each of the conserved motifs, we created a profile HMM (pHMM) signature generalizing the motif as described in detail in the Methods section. Next, we identified all NB-ARC and NACHT domain proteins encoded by genes in the genomic vicinity (within 5 kbp) of the genes encoding our set of Bell-domain proteins. Then, we analyzed N-termini of of these NLR-like proteins for occurrences of the pHMM-signatures of Bell-associated C-terminal motifs. We found that half of the motifs identified at least one gene pair encoding a Bell-domain protein and a NACHT/NB-ARC protein. To be more stringent in our analyses, we focused further investigations only on motifs that identified at least 5 non-redundant gene pairs which reduced our set to 29 motifs (comprising up to 75 non-redundant gene pairs per motif) (Table S2). The motifs were clustered based on the overlap between the matched gene pairs (Fig. S2). When at least half of the gene pairs identified for a given motif were also matched by another motif, the two motifs were considered related and were grouped in the same motif cluster. The procedure led to identification of nine clusters (some of which comprised in fact a single motif) (Fig. S2). In each cluster, we selected a representative motif on the basis of the number of underlying sequences (Fig. 2). The motifs families defined by each cluster were termed BASS families (for bacterial amyloid signaling sequence) (Fig. 2, Fig. S2). As expected, we recovered the already identified RHIM-related motif represented by 19 gene pairs (BASS3). We also carried out an inverse screen in which we looked for conserved gapless motifs in N-terminal sequences of the NB-ARC/NACHT protein set, generated pHMM signatures for the extracted motifs, and searched for their occurrences in C-termini of the Bell proteins. We retained 12 motifs identifying at least five non-redundant gene pairs and clustered these motifs within the previously defined clusters. While eleven grouped to one of the nine previously identified families (Fig. S2), one novel motif did not, and is thus defining an additional family, we term BASS10.

**Figure 2.**
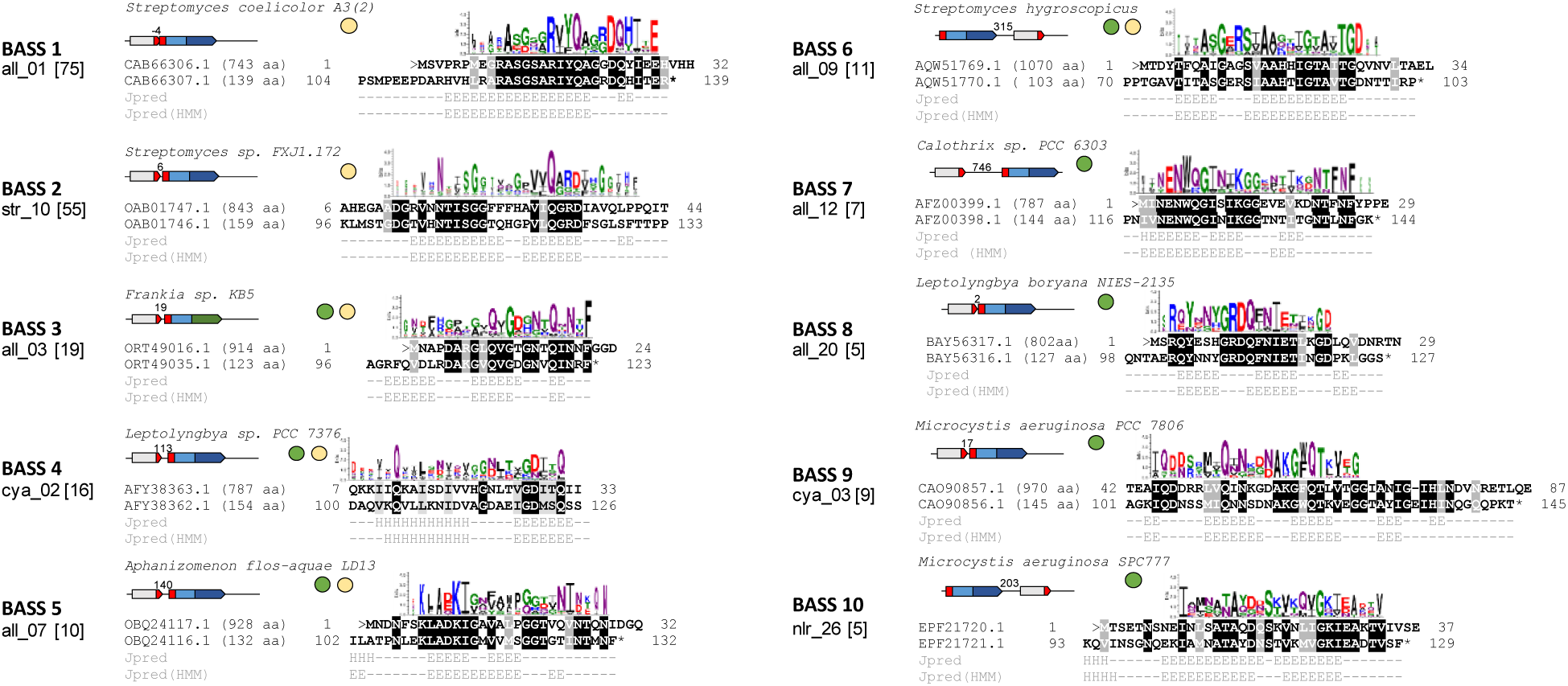
Ten bacterial amyloid signaling sequences motifs (BASS). For each of the ten identified motifs, a representative gene pair is given. A consensus sequence for the motif is given. The consensus was generated using Weblogo from the alignment of all motifs pairs bearing the corresponding motif. For each gene pair chosen as illustrative example, the species name, the genome architecture, the gene number and protein size are given as well as an alignment of the Bell-domain and NLR-associate motifs. The number given above the gene diagram is the distance between the Bell-domain encoding and NLR encoding ORF, negative number represent gene overlaps. The sequences encoding the BASS motif is represented in red, the Bell-domain in grey, NB-ARC domain is light blue, TPR repeats in dark blue and WD repeats in green. Under the alignment the secondary structure prediction for the individual sequences of the Bell-domain motif (Jpred) or for a HMM-alignment of the sequence (Jpred (HMM)) are given (E for extended, H for helical). The number of motif pairs identified with the different motifs is given in parentheses. The dots symbolize the phylogenetic distribution of the motif (green, cyanobacteria, beige, actinobacteria).

We complemented the search with an alternative procedure consisting of the local pairwise alignment of adjacent Bell C-termini and NLR N-termini. Highly conserved pairs of a length of at least 15 amino acids were found in 283 gene pairs including 44 pairs not matched previously with the BASS1 to 10 motifs. In this number were two pairs from archaeal Methanosarcinales species (Table S2).

We analyzed the distance and the relative orientation between the Bell encoding gene and the adjacent NLR-encoding gene in our set of 346 matching gene pairs (Table S2). Genes were collinear in 91.3% of the cases. In most collinear gene pairs, the distance between the genes was very small, in 57% of the cases the distance between the two ORFs was 10 bp or less and in 42% of the cases the two ORFs overlapped. In one extreme case, in *Spirosome oryzae*, the Bell domain encoding gene and the NLR-encoding gene overlapped by 57 bp. Gene overlaps are not uncommon in prokaryotes, roughly 20% of the collinear gene pairs overlap, (Fukuda et al. 2003). It is likely that the Bell and NLR-encoding genes pairs with short intergenic distances or gene overlaps are transcribed as polycistronic messenger RNAs reflecting the possible functional link and/or co-regulation between the genes in the pair.

### Paired BASS motifs occur in multicellular species

Mirroring the distribution of Bell-domains and NLRs, the vast majority of BASS-motifs pairs occur in *Actinobacteria* and *Cyanobacteria* (Table S1) with the filamentous genus *Streptomyces* dominating the data set with 105 pairs (Table S2). Some motifs appear specific to *Actinobacteria* (BASS 1, 2) others to *Cyanobacteria* (BASS 7, 8, 9, 10), others occur in both classes (BASS 3, 4, 5, 6), (Table S2). All actinobacterial hits occur in filamentous species with the possible exception of *Arthrobacter sp. Rue61a*. No direct information on the growth morphology for this specific strain could be recovered but the corresponding genus is described as unicellular. The hits in *Cyanobacteria* correspond essentially to multicellular filamentous species (*Nostoc, Pseudoanabaena, Calothrix, Aulosira, Tolothrix, Leptolyngbya* genera). Some species with hits (*Microcystis aeruginosa* and *Crocosphaera watsonii*) are described as unicellular (non-filamentous) *Cyanobacteria* but are also found in multicellular aggregates. One hit was found in *Cyanothece* sp. PCC 7424 which is a unicellular strain reported to form large amounts of mucilage, as well as in *Chamaesiphon minutus* PCC 6605 a unicellular species forming epiphytic colonies on aquatic plants. *Cyanothece, Microcystis, Crocosphaera* and Chamaesiphon are however distinct from the main group of unicellular Cyanobacteria (*Prochlorococcus, Synechococcus*) in terms of phylogenetic position and abundance of secondary metabolism clusters and homologs of *AmiC* involved in multicellular growth (Shih et al. 2013). The remaining gene pairs occur in *Archaea, Bacteriodetes* and *Chloroflexi,* in multicellular species, which are either filamentous (like the Chloroflexi *Ktedonobacter racemifer*) or existing as cellular aggregates (like the *Archaea Methanosarcinales* species). We conclude that matching BASS-motif pairs occur with a wide phylogenetic distribution but in the vast majority of cases in multicellular strains and genera.

### Sequence analysis of BASS motifs

BASS motifs were typically around 25 amino acid in length (Fig. 2). All motifs have a predicted propensity for β-sheet formation (of note is the fact that this criterion was not used for their identification). The sequence logos of the motifs show conservation of G, N and Q residues as well as charged residues and conserved patterns of hydrophobic residues. In comparison to the total of Uniprot entries, the motifs are enriched for Q, N, G, V and I and depleted in K, P and L (Table S3). Kajava and co-workers have reported a similar bias in amino acid composition of β-helical proteins with enrichment in N, V and G and depletion in P and L (Baxa et al. 2006). Fungal amyloid motifs of the HRAM, sigma and PP family share a similar bias in composition, residue conservation and length (Daskalov et al. 2012) (Table S3).

Algorithms predicting amyloid propensity derived from the analysis of pathological amyloids generally perform poorly when run on fungal functional amyloid motifs (Ahmed and Kajava 2013). In contrast, the ArchCandy program predicting propensity to form β-arch structures shows good performance for pathological amyloids but also in the case of fungal functional amyloid motifs (Ahmed et al. 2014). We thus evaluated the amyloid propensity of BASS motif containing proteins with ArchCandy and found that their amyloid propensity scores were comparable to those calculated for a selection of validated fungal amyloid motifs and human RHIM-motifs (Fig. S3) (BASS1 and BASS3 scored below the recommended stringent score threshold (0.575) but this was also the case for an HRAM5 fungal prion motif (Daskalov et al. 2015a)). In each case, the high scoring region matched the position of the identified BASS motif. Based on this analysis, the identified bacterial motifs are predicted to have amyloid forming propensity (note again that they were not selected on this criterion).

The fungal HRAM motifs exemplified by the HET-s prion-forming domain present a two-fold pseudorepeat of the motif in the effector protein (Daskalov et al. 2015b; Ritter et al. 2005). The two pseudorepeats are separated by a variable flexible glycine and proline -rich loop of 6-15 amino acids in length (Daskalov et al. 2015a; Wasmer et al. 2008). We find that some BASS motifs also occur as pseudorepeats. In proteins pairs of different *Streptomyces* species, BASS1 and 6 motifs are present in the Bell-domain protein as two (or three) pseudorepeats separated by a variable proline-rich region and as a single repeat in the corresponding NLR (Fig. S4). These motifs thus are similar in this respect to fungal HRAM motifs.

Fungal amyloid signaling motifs can be associated to other effector domains not related to the HeLo-family (HeLo, Helo-like, sesA) but equally encoded by NLR/effector gene pairs (Daskalov et al. 2016; Daskalov et al. 2012; Dyrka et al. 2014). These domains include α/β-hydrolase and PNP-UDP phosphorylase domains. We wondered therefore whether BASS motifs could also be associated to domains distinct from Bell. Thus, we analyzed the occurrence of identified BASS motifs in gene pairs encoding a NLR and a second protein, for which the motif is found in the C-terminal region next to a domain distinct from Bell. At least four such situations were identified with BASS motifs associated to the following domains : a α/β-hydrolase domain, the TIR2 domain, the CHAT metacaspase domain and a guanylate cyclase domain (Fig. S5). All identified examples were found in multicellular species. Bacterial genomes contain genes in which the same effector domains are found in an “all-in-one” association, where the effector domain is directly associated to the NOD and repeat domains of the NLR-like protein, as describe previously for fungal NLRs (Daskalov et al. 2012). While some of these domains have rather general functions, it is of interest to note that all these domains (α/β-hydrolase, TIR2, CHAT and guanylate cyclase) are to various extends documented to be involved in immune and programmed cell death pathways (Daskalov et al. 2016; Freihat et al. 2019; Koonin and Aravind 2002; Nimma et al. 2017; O’Neill and Bowie 2007; Xue et al. 2012).

### BASS1 and BASS3 from selected bacterial species behave as prion-forming domains when expressed in Podospora

NLR-associated fungal amyloid motifs were initially identified as prion-forming domains (Daskalov et al. 2012). In order to determine if the BASS motifs could behave similarly, we expressed selected BASS motifs in *P. anserina*. used to analyze prion propagation of both homologous and heterologous amyloid signaling motifs (Benkemoun et al. 2011; Daskalov et al. 2015b; Daskalov et al. 2016). It is a valuable alternate model to yeast which was also extensively used to document prion properties of heterologous sequences. In *P. anserina* prion propagation and transmission are easy to monitor because of the syncytial structure of the mycelium and because strains spontaneously fuse and mix their cytoplasmic content when confronted (Benkemoun et al. 2006). We chose a motif corresponding to the most populated family (BASS 1) from the model species *Streptomyces coelicolor* A3(2) and three phylogenetically diverse BASS3 (RHIM-like motifs) form *Actinobacteria* species *Streptomyces atratus* and *Nocardia fusca* and form the *Cyanobacteria* species *Nostoc punctiforme* (Fig.2, Fig.S6, Table 3). The three selected BASS 3 sequences were diverse and share <50% identity between species (Fig.S6). We expressed both the Bell-associated BASS and the corresponding NLR-associated BASS (except for *N. fusca* proteins for which only the Bell-associated sequence was studied) (Table 3). The different BASS motifs were fused with either GFP (in N-terminus) or RFP (in C-terminus) and expressed under a strong constitutive promotor in *P. anserina*. All fusion proteins showed bistability, subsisting initially in a diffuse cytoplasmic state that could convert over time to discrete cytoplasmic dots as previously described for fungal prion-forming domains (Table 4, Fig.3) (Balguerie et al. 2003; Benkemoun et al. 2011; Coustou-Linares et al. 2001; Daskalov et al. 2015b; Daskalov et al. 2016). By analogy with fungal prions, we denote the diffuse state [b*] and the aggregated state [b]. Transition from the diffuse [b*] to the aggregated [b] state could occur either spontaneously or be induced by cytoplasmic contact with a strain expressing the fusion protein in [b] state (Fig.4). We conclude that all tested BASS motifs direct infectious propagation of the aggregated state and thus behave as prion-forming domains in this fungal model.

**Figure 3.**
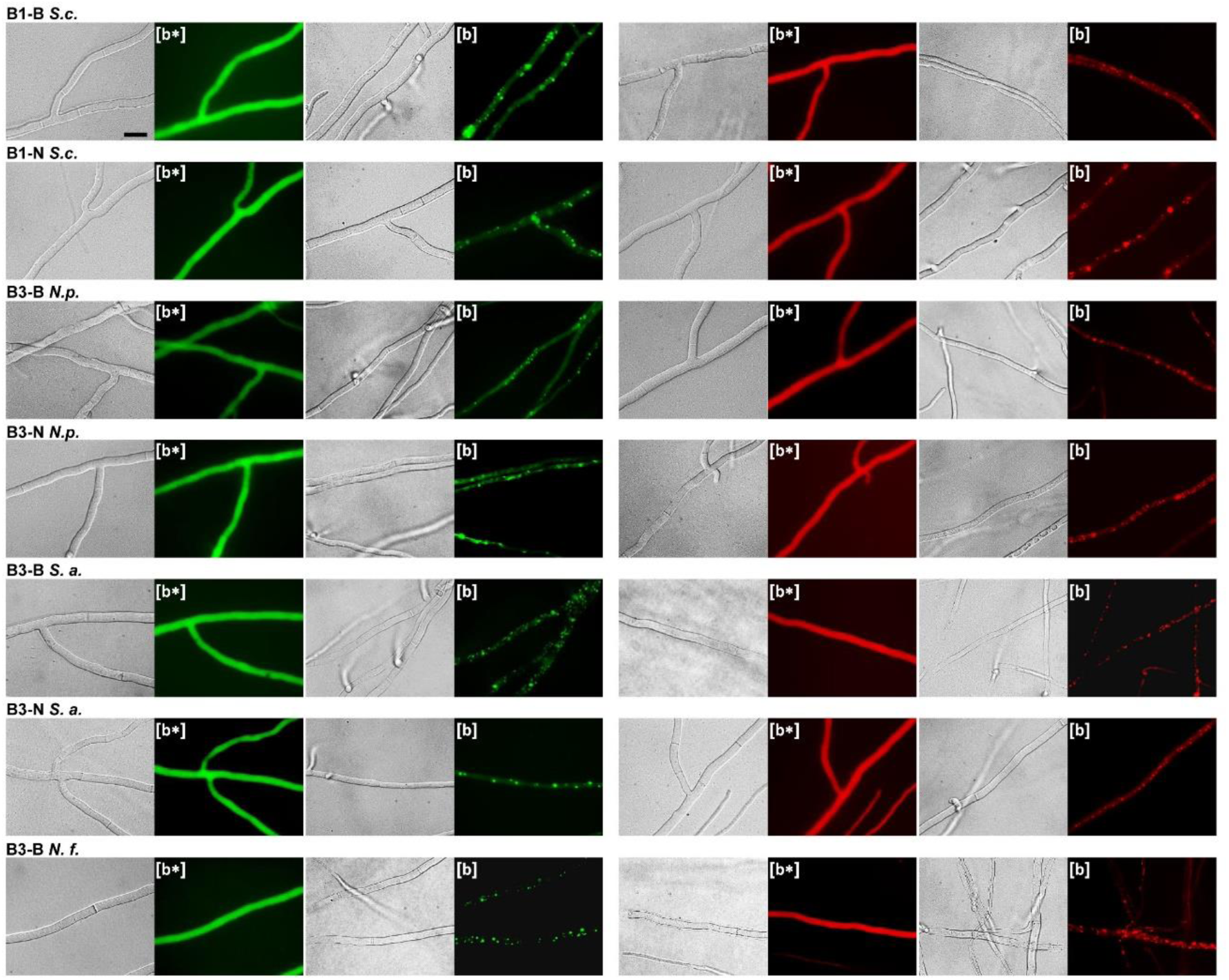
BASS-motif prion a formation in *Podospora anserina*. Micrographs of *P. anserina* strains expressing molecular fusions of BASS1 and BASS3 motifs with GFP (in N-terminus) or RFP (in C-terminus) as indicated above each micrograph (Scale bar 5 μm). Transformants displayed an initial diffuse fluorescence noted [b*] phenotype (left side of the panels) and acquired dot like fluorescent aggregates, [b] phenotype, after contact with strains already spontaneously expressing the infectious aggregated [b] state (right side of the panels).

**Figure 4.**
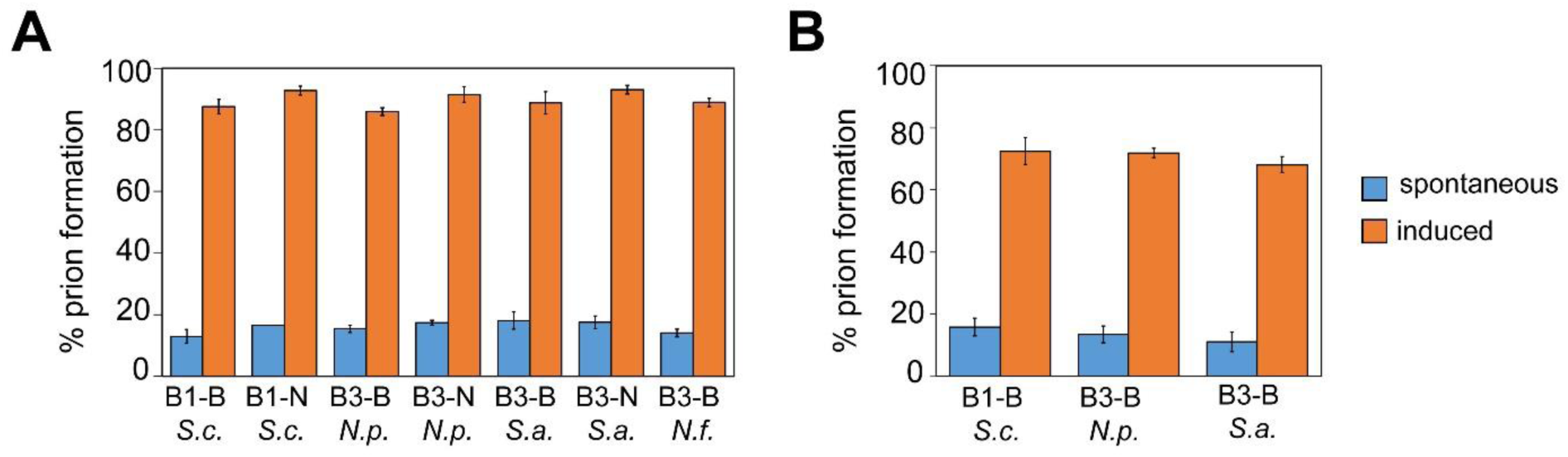
BASS motifs propagate as prions in *Podospora anserina* and NLR-side motifs induce prion formation of Bell-side motifs. **A.** Histogram representing the percentage of [b*] strains expressing the given BASS motifs (fused to RFP in C-terminus) converted to prion-infected [b] phenotype after contact either with non-transfected prion-free control strain (spontaneous, in blue), or with prion [b] strains expressing the same motif in the aggregated state (induced, in orange). Percentages are expressed as the mean value ± standard deviation for three independent experiments using six subcultures of four different transformants and corresponding to the analysis of ∼70 subcultures for each BASS motif. In each case, the BASS motif are fused to RFP. **B.** Histogram representing the percentage of [b*] strains expressing the given BASS motif (fused to RFP in C-terminus) as indicated converted to prion-infected [b] phenotype by contact either with non-transfected prion-free control strains (spontaneous, in blue), or with prion [b] strains expressing the corresponding NLR-side motif fused to GFP (in N-terminus) in the aggregated states (induced, in orange). Percentages are expressed as the mean value ± standard deviation for experiments using six to twelve subcultures of six to ten different transformants and correspond to the analysis of 80 to 100 subcultures for each BASS motif.

**Figure 5.**
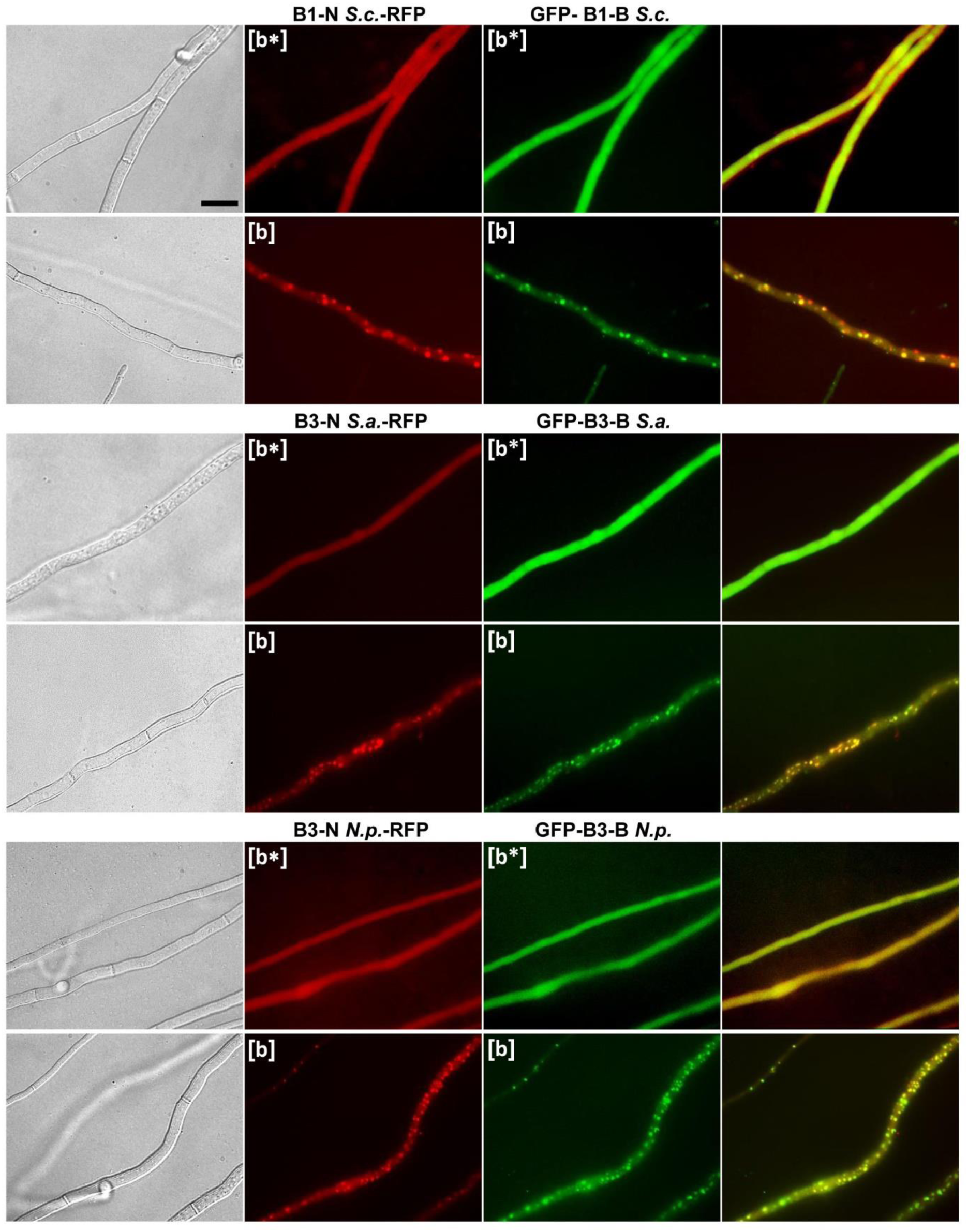
Bell and NLR-side BASS motif co-aggregation in *Podospora anserina*. Micrographs of *Podospora anserina* strains co-expressing Bell-side BASS motifs fused to GFP (in N-terminus) as indicated and the corresponding NLR-side motif fused to GFP (in C-terminus), (Scale bar 5 μm). Panel are from left to right, bright field, RFP, GFP and overlay.

**Figure 6.**
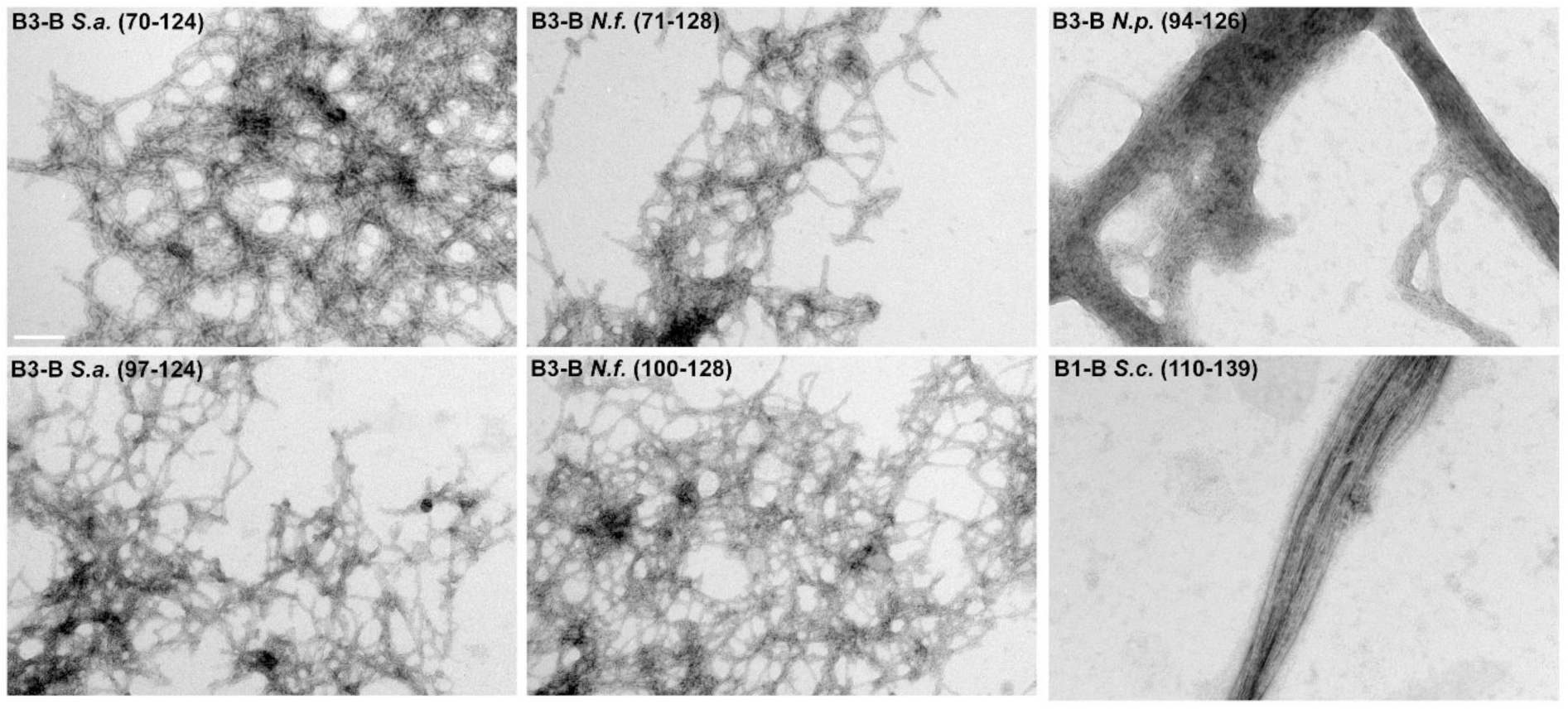
Electron microscopy of BASS-motif fibrils. Electron micrographs of fibrils formed by selected BASS motifs as indicate (scale bar, 100nm).

**Table 3.** Sequences selected for functional studies.

| Motif/Domain/Strain | Accession number and coordinates | Code name |
| --- | --- | --- |
| BASS1/Bell/*Streptomyces coelicolor* A3(2) | CAB66307.1 (110-139) | B1-B *S.c.* |
| BASS1/NLR/*Streptomyces coelicolor* A3(2) | CAB66306.1 (1-34) | B1-N *S.c.* |
| BASS3/Bell/*Streptomyces atratus* | WP_037701008.1 (70-124) | B3-B *S.a.* |
| BASS3/NLR/*Streptomyces atratus* | WP_037701012.1 (1-37) | B3-N *S.a.* |
| BASS3/Bell/*Nostoc punctiforme* PCC 73102 | ACC79696.1 (94-126) | B3-B *N.p.* |
| BASS3/NLR/*Nostoc punctiforme* PCC 73102 | ACC79697.1 (1-38) | B3-N *N.p.* |
| BASS3/Bell/*Nocardia fusca* | WP_063130184.1 (74-128) | B3-B *N.f.* |

**Table 4.**
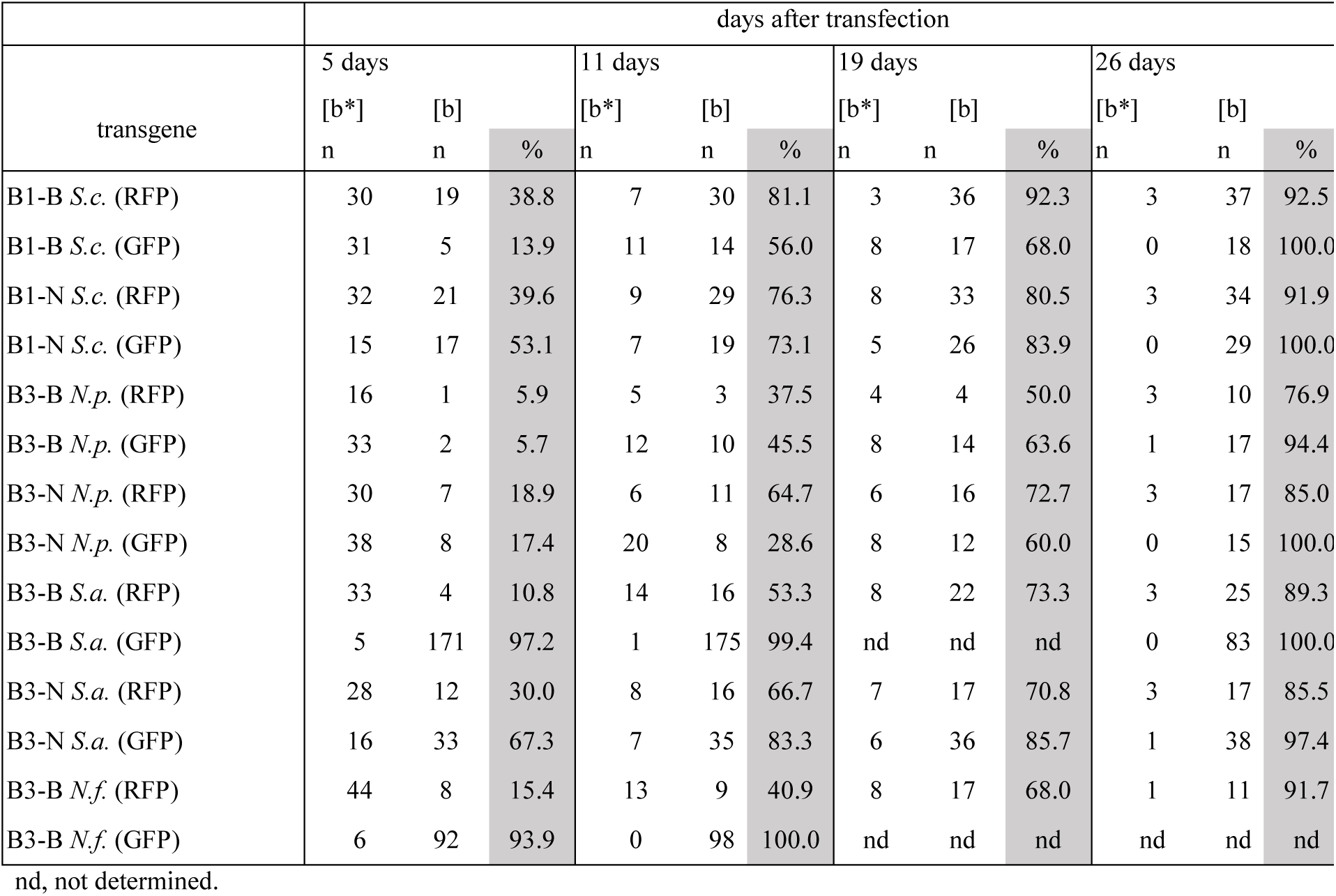
Spontaneous prion formation rates of selected bacterial motifs.

| transgene | days after transfection |  |  |  |  |  |  |  |  |  |  |  |
| --- | --- | --- | --- | --- | --- | --- | --- | --- | --- | --- | --- | --- |
|  | 5 days |  |  | 11 days |  |  | 19 days |  |  | 26 days |  |  |
|  | [b*] | [b] |  | [b*] | [b] |  | [b*] | [b] |  | [b*] | [b] |  |
|  | n | n | % | n | n | % | n | n | % | n | n | % |
| B1-B *S.c.* (RFP) | 30 | 19 | 38.8 | 7 | 30 | 81.1 | 3 | 36 | 92.3 | 3 | 37 | 92.5 |
| B1-B *S.c.* (GFP) | 31 | 5 | 13.9 | 11 | 14 | 56.0 | 8 | 17 | 68.0 | 0 | 18 | 100.0 |
| B1-N *S.c.* (RFP) | 32 | 21 | 39.6 | 9 | 29 | 76.3 | 8 | 33 | 80.5 | 3 | 34 | 91.9 |
| B1-N *S.c.* (GFP) | 15 | 17 | 53.1 | 7 | 19 | 73.1 | 5 | 26 | 83.9 | 0 | 29 | 100.0 |
| B3-B *N.p.* (RFP) | 16 | 1 | 5.9 | 5 | 3 | 37.5 | 4 | 4 | 50.0 | 3 | 10 | 76.9 |
| B3-B *N.p.* (GFP) | 33 | 2 | 5.7 | 12 | 10 | 45.5 | 8 | 14 | 63.6 | 1 | 17 | 94.4 |
| B3-N *N.p.* (RFP) | 30 | 7 | 18.9 | 6 | 11 | 64.7 | 6 | 16 | 72.7 | 3 | 17 | 85.0 |
| B3-N *N.p.* (GFP) | 38 | 8 | 17.4 | 20 | 8 | 28.6 | 8 | 12 | 60.0 | 0 | 15 | 100.0 |
| B3-B *S.a.* (RFP) | 33 | 4 | 10.8 | 14 | 16 | 53.3 | 8 | 22 | 73.3 | 3 | 25 | 89.3 |
| B3-B *S.a.* (GFP) | 5 | 171 | 97.2 | 1 | 175 | 99.4 | nd | nd | nd | 0 | 83 | 100.0 |
| B3-N *S.a.* (RFP) | 28 | 12 | 30.0 | 8 | 16 | 66.7 | 7 | 17 | 70.8 | 3 | 17 | 85.5 |
| B3-N *S.a.* (GFP) | 16 | 33 | 67.3 | 7 | 35 | 83.3 | 6 | 36 | 85.7 | 1 | 38 | 97.4 |
| B3-B *N.f.* (RFP) | 44 | 8 | 15.4 | 13 | 9 | 40.9 | 8 | 17 | 68.0 | 1 | 11 | 91.7 |
| B3-B *N.f.* (GFP) | 6 | 92 | 93.9 | 0 | 98 | 100.0 | nd | nd | nd | nd | nd | nd |

nd, not determined.

### NLR-side BASS induce prion conversion of Bell-side BASS

If BASS motifs are analogous to fungal amyloid signaling motifs, it is expected that the NLR-side and Bell-side BASS co-aggregate and that an aggregated NLR-side BASS is able to convert the matching Bell-side BASS to the prion state (Daskalov et al. 2015b). To test whether BASS motif pairs could co-aggregate, we co-expressed each motif pair in the same fungal strain. We found that all tested pairs co-localize in dots although in some cases the co-localization was not complete (Fig. 5, Fig. S7). Then, we tested whether the NLR-side BASS in the [b] prion state could induce prion aggregation of the corresponding Bell-side BASS. Strains expressing RFP-fused Bell-side motifs in the non-prion [b*] state were confronted with strains expressing the corresponding GFP–fused NLR-side motifs in the [b] prion state. In all tested cases, we found that the NLR-side BASS efficiently converts the Bell-side BASS to the prion state (Fig. 6). The level of amino acid identity within cross-seeding BASS motif pairs is in the range of 43-56% (Fig. 2, Fig. S6) and comparable to the level of identity leading to prion cross-seeding of fungal amyloid signaling motifs (Benkemoun et al. 2011; Daskalov et al. 2015b; Daskalov et al. 2016). When expressed in this fungal model, tested BASS are functionally analogous to fungal amyloid signal motifs in the sense that the NLR-side motif is able to interact with the Bell-side motif and to induce its prion aggregation.

### BASS1 and BASS3 form fibrils *in vitro*

To verify the prediction that the identified bacterial motifs correspond to amyloid forming sequences, we produced the selected Bell-side BASS1 motif (*Streptomyces coelicolor* A3(2)) and the three BASS3 motifs (*Streptomyces atratus, Nocardia fusca* and *Nostoc punctiforme*) in recombinant form and analyzed their ability to form fibrils *in vitro*. The proteins encompassing the motifs were expressed as inclusion bodies, purified under denaturing conditions in 8M urea. Upon dilution of the denaturant, all constructs led to the formation of fibrillar aggregates, either as dispersed fibrils or as laterally associated large bundles resembling those formed by HET-s(218-289) (Balguerie et al. 2003; Sabate et al. 2007). Fibril width was in the range of 5-7 nm comparable to the previously identified fungal signaling motifs (Fig. 6), (Daskalov et al. 2015b; Daskalov et al. 2016; Sabate et al. 2007). We conclude that the selected BASS motifs spontaneously assemble into fibrils *in vitro*.

For the *Streptomyces coelicolor* A3(2) BASS1 motif (CAB66307.1, residue 38 to 139), belonging to the most commonly occurring BASS family, we further analyzed structural properties of the *in vitro* fibrils (Fig. 7A). We employed X-ray fiber diffraction to examine the presence of a cross-β architecture in the sample. We observed an intense ring at 4.7 Å, and a weak ring at 10 Å (Fig. 7B), this pattern being characteristic for a cross-β structure corresponding to the inter-strand and inter-sheet spacing respectively, typically observed in amyloid fibrils (Sunde et al. 1997). Next, we produced a BASS1 sample isotopically and uniformly ^13^C labeled to carry out solid-state NMR analysis. Cross-polarization ^13^C experiment (Fig. 7C) revealed a well-resolved spectrum, implying a high structural order at the local level (Loquet et al. 2018a). In line with the X-ray diffraction analysis, solid-state NMR ^13^C chemical shifts of BASS1 indicate a protein conformation rich in β-sheet secondary structure, illustrated with a high field effect of the carbonyl region indicative of β-sheet structure (Wang and Jardetzky 2002). Taken together, these analyses show that this BASS-motif forms β-sheet-rich amyloid fibrils, with NMR features highly comparable to amyloid fibrils of the fungal HET-s(218-289) (Ritter et al. 2005; Siemer et al. 2005).

**Figure 7:**
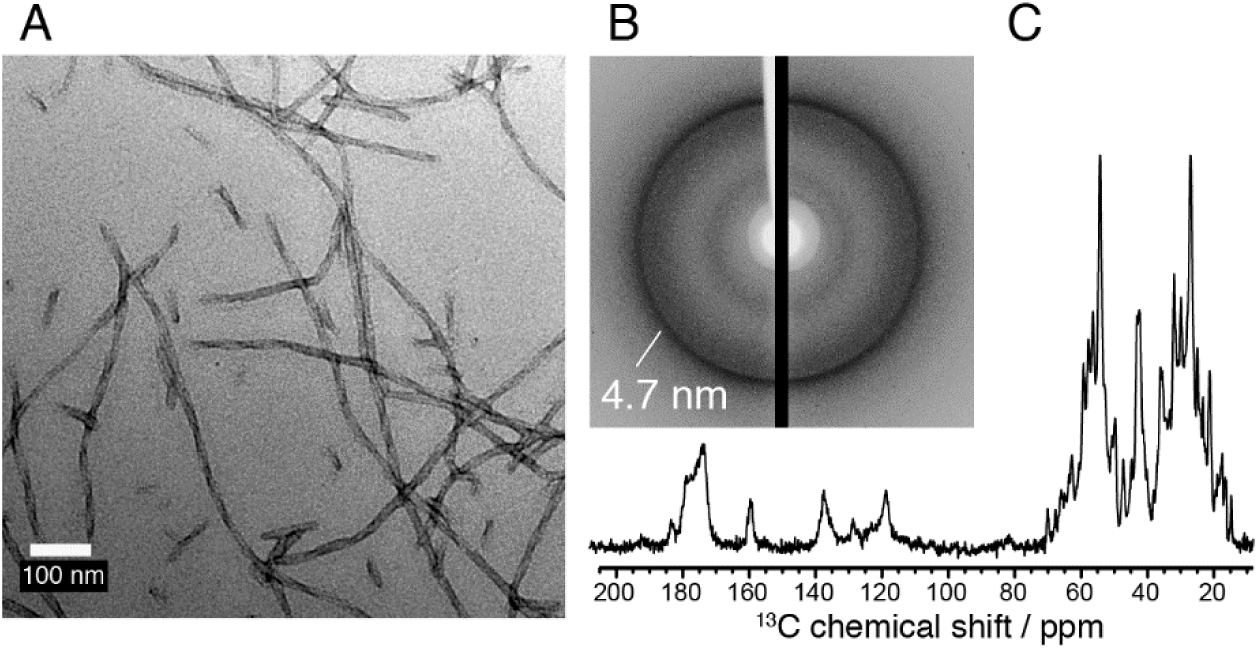
Structural analysis of the *S. coelicolor* BASS 1 motif. **A.** Negatively-stained electron micrograph of BASS 1 fibrils (B1-B S.c.(38-139), CAB66307.1), scale bar is 2 µm. **B.** X-ray diffraction pattern of BASS 1 fibrils, reflection at 4.7 Å is highlighted, corresponding to the inter-strand spacing. **C.** ^13^C solid-state NMR spectrum of BASS 1 fibrils.

## Discussion

Amyloids have initially been identified in the context of human protein-deposition diseases and correspond to protein aggregates with a cross-β structure (Riek and Eisenberg 2016). The nucleated-polymerization process governing their assembly allows some amyloids to propagate their conformational state as prions (Colby and Prusiner 2011; Wickner et al. 2016). The amyloid fold also plays a variety of functional roles (Loquet et al. 2018b; Otzen and Riek 2019). In particular, functional amyloids have been found to be involved in signal transduction cascades controlling programmed cell death processes. In mammals, the RHIM motif controls assembly of the RIPK1/RIPK3 complex in the necroptosis pathway (Li et al. 2012; Mompean et al. 2018). In fungi, amyloid motifs control a signal transduction mechanism based on transmission of an amyloid fold from an NLR protein to downstream cell-death execution proteins (Daskalov et al. 2015b; Daskalov et al. 2016). In the prokaryotic reign, functional amyloids have been found to be involved in biofilm formation, development and virulence (Erskine et al. 2018; Rouse et al. 2018; Van Gerven et al. 2018). In addition, prion formation has been reported in bacterial model systems (Giraldo et al. 2011; Shahnawaz et al. 2017; Wang et al. 2017). We show here that phylogenetically diverse bacterial lineages including *Actinobacteria* and *Cyanobacteria* display fungal-like amyloid signaling motifs and thus suggest that NLR-associated amyloid signaling is also present in these multicellular bacterial lineages.

### Diversity of amyloid signaling motifs in Bacteria

Several families of amyloid signaling motifs were described in fungi and remarkably, PP, one such family appears to be related to the animal RHIM amyloid motifs (Daskalov et al. 2016; Li et al. 2012). We now find that a motif similar to RHIM/PP and occurring in the same domain architectures as the fungal PP-motif also exists in Bacteria. Although, the motif consensus is different for RHIM, PP and BASS3 motifs, all share the central G-φ-Q-φ-G signature (Fig. S6). This signature corresponds to the central core of the RHIM amyloid structure with the tight interdigitation of two such motifs in one β-strand layer (Mompean et al. 2018). The extension of the presence of this amyloid motif to prokaryotic lineages supports the hypothesis of long-term conservation of this motif for amyloid signaling purposes from bacteria, to metazoan and fungi. However, due to the moderate sequence similarity between the motifs the possibility of a convergent evolution towards the amyloid signaling function cannot be ruled out.

We have also identified nine other bacterial amyloid motif families. This finding suggests an extensive diversification of this type of signaling domains in prokaryotes. In fungi, amyloid signaling motifs were also found to be diverse with so far three main families described (HRAM, PP and sigma) (Daskalov et al. 2015a; Daskalov et al. 2012). Diversity of such motifs in bacterial lineages appears even greater that in fungi, which might be expected considering the larger phylogenetic breadth of the bacterial lineages compared to fungi (Hug et al. 2016). Diversity of bacterial motifs almost certainly exceed the 10 families described herein, considering that we restricted the analysis to motifs recovering at least 5 non-redundant matching gene pairs. Except for the RHIM/PP-motifs there is no obvious sequence homology between bacterial and fungal motifs, which is not surprising considering that some of the BASS motifs appear specific for a given bacterial phylum. In spite of the lack of direct sequence homology, bacterial and fungal motifs have common features, they show a similar length (typical 20-25 amino acids) and amino acid composition biases, apparently typical of proteins forming β-arch structures (Baxa et al. 2006). In addition, some bacterial motifs appear as double (or triple) pseudo-repeats as described in the case of fungal HRAMs (Daskalov et al. 2015a; Ritter et al. 2005).

### BASS form prions

When expressed in *Podospora anserina*, selected BASS motifs behaved as prion-forming domains and thus were functionally analogous to previously characterized fungal prion signaling motifs. In other words, they are capable, *in vivo*, in this heterologous setting, to form aggregates and to propagate this aggregation state as prions. In addition, we find that the NLR-side and Bell-side of matching motifs are able to co-aggregate and that, NLR-side motifs convert Bell-side motifs to the prion state. These results are consistent with the proposition that the motifs functionally interact and that the NLR-side motif serves as a template for the transconformation of the Bell-side motif as shown for the fungal prion signaling motifs (Daskalov et al. 2015b). We find that the same motifs form fibrils *in vitro*. In case of the BASS1 motif of *Streptomyces coelicolor,* X-ray diffraction and solid-state NMR analyses indicate the formation of a highly-ordered cross-β structure. Comparable observations have been made using the same biophysical techniques for analogous fungal prion-forming domains such as HET-s(218-289) (Ritter et al. 2005; Siemer et al. 2005; Wan and Stubbs 2014), NWD2(1-30) (Daskalov et al. 2015b) or PP (Daskalov et al. 2016). It thus appears that for the BASS motifs that were tested experimentally, we confirm the identified sequences are amyloid prions, again making is plausible that the other motifs also represent analogous prion amyloids.

### NLRs, Bell domains and amyloid signaling in multicellular bacteria

Proteins with NLR domain architectures control various biotic interactions in plants, animals and fungi (Jones et al. 2016; Uehling et al. 2017). Throughout, we have used the designation NLR to specify proteins displaying a domain architecture associating a NB-ARC or NACHT-type NOD domain (nucleotide binding and oligomerization domain) and ANK, TPR, WD or LRR super-structure forming repeats (SSFR). While some authors reserve the NLR designation to plant and animal NBS-LRR proteins, we adhere to an extended assertion of the term also including NB-SSFR proteins found in fungi and prokaryotes. It has been reported previously that bacterial genomes (in particular in *Actinobacteria* and *Cyanobacteria*) display genes encoding proteins with a NBS-SSFR architecture (Asplund-Samuelsson et al. 2012; Koonin and Aravind 2002; Urbach and Ausubel 2017). In fungi, both NACHT/WD and NB-ARC/TPR proteins have been shown to control non-self recognition and programmed cell death in different species (Chevanne et al. 2009; Choi et al. 2012; Espagne et al. 2002; Heller et al. 2018; Saupe et al. 1995). Urbach and Ausubel have shown that these NACHT/WD and NB-ARC/TPR architectures are ancestral to the NBS/LRR architecture, which represent a more recent acquisition (that occurred independently in plant and animal lineages).

In our survey of over 100 000 prokaryotic genomes, we find again that NLR architecture proteins are characteristic of bacterial lineages encompassing genera and species with a multicellular organization (filamentous or multicellular aggregates forming species). These are found in filamentous *Actinobacteria*, *Cyanobacteria* and *Chloroflexi* and also in aggregate-forming Archaea. Characteristic of those bacterial lineages that display NLRs is also the presence of the Bell domain. Like fungal NLRs, prokaryotic NLRs typically have NB-ARC/TPR or NACHT/WD architectures and do not display LRR repeats (Dyrka et al. 2014; Koonin and Aravind 2000; Urbach and Ausubel 2017). Another communality between fungal and bacterial NLRs is the presence in both lineages of NLR with WD and TPR-repeats showing high levels of internal conservation (with repeat units showing 80-90% sequence identity, a situation in stark contrast with the bulk of the WD and TPR-repeats in proteins), (Dyrka et al. 2014), (Hu et al. 2017; Marold et al. 2015). In fungi, this internal conservation is related to a mechanism of rapid diversification of repeat domain binding specificity (Paoletti et al. 2007). These NLR-repeats are under positive Darwinian selection and diversify rapidly by a mechanism of repeat-unit reshuffling representing both the cause and consequence of their high internal conservation (Chevanne et al. 2010; Dyrka et al. 2014; Paoletti et al. 2007). Analogous WD and TPR repeats in bacterial NLRs might similarly be under a specific evolutionary regimen allowing rapid diversification. The possibility of an ancestral role of NLR-like proteins in programmed cell death and host defense in multicellular bacteria has been raised before (Koonin and Aravind 2002). Together with the communalities mentioned above, the mechanistic similarity between fungal NLR/amyloid motif/HeLo and bacterial NLR/amyloid motif/Bell domain associations we report, now makes it indeed plausible to envision that the bacterial gene pairs equally function in host defense and programmed cell death. In this hypothesis, the Bell domain might correspond to a programmed cell-death execution domain. In support of this view is the sequence similarity between the signature consensus sequences in the N-terminal α-helical region of Bell, fungal HeLo, metazoan MLKL and plant CC-domains (Fig. 1). Functional studies of the Bell-domain and its possible function in cell death execution are now required to explore this hypothesis. Of note is also the fact that other protein domains related to immune functions and programmed cell death in animals and plants (such as the TIR and caspase-like CHAT domain) can also be found associated to amyloid signaling motifs and that the same domains are frequently found as N-terminal domain of NLRs in multicellular bacteria (Table S1).

Remarkably, in fungi NLRs are restricted to filamentous genera (both in Ascomycota and Basidiomycota) and are not found in yeast species (Dyrka et al. 2014). This situation is mirrored in the present study by the fact that NLRs in general as well as the matching NLR/Bell gene pairs are found in *Actinobacteria* and *Cyanobacteria* species with a multicellular morphology but appear to be absent or rare in phylogenetically related unicellular classes. Although, PCD pathways also exist in unicellular species those continue to represent a paradox (Durand et al. 2016; Koonin and Krupovic 2019). Defense-related PCD is generally considered an attribute of multicellular organisms in which altruistic cell death can be advantageous for the survival of the multicellular organism (Iranzo et al. 2014). Our results suggest that filamentous fungi and bacteria have in common the use of NLR-associated amyloid signaling processes. We propose that this shared molecular mechanistic feature between filamentous fungi and filamentous *Actino* and *Cyanobacteria* stems from their common cellular organization and life style. These fungi and bacteria are phylogenetically distant but morphologically alike. In a similar way, abundance and complexity of secondary metabolism clusters involved in allelopathic interactions appears to be a specific feature of filamentous fungi and bacteria compared to related unicellular genera (Keller 2019; Shih et al. 2013; van der Meij et al. 2017). Presence of NLRs and amyloid signaling might represent a common genome hallmark of diverse multicellular microbes. As multicellular organisms, microbes from both reigns could have to cope with parasites and pathogens and thus might rely on altruistic cell suicide for defense, as an immune-related programmed cell death mechanism akin to those operating in multicellular plant and animals lineages. We propose based on these genomic and functional analogies that filamentous fungi and filamentous bacteria share in common the use of NLR-associated amyloid motifs in the control of immune-related programmed cell death. The implication of this hypothesis is that the use of NLRs for immune-related functions might be ancestral and shared universally between multicellular Archaea, Bacteria, fungi, plants and animals. The mechanistic resemblance we report between prokaryotic and eukaryotic NLRs, invites to reconsider the evolutionary trajectory of this protein family, which might have a very ancient history in the control of biotic interactions.

## Methods

### Homology searches

The N-terminal 102 amino acid-long fragment of protein ONI86675.1 (A0A1V2QF20_9PSEU) (Yeager et al. 2017) was used as the Bell domain query in Jackhmmer (Eddy 2011; Potter et al. 2018) (20 iterations, 2133 hits) and Psi-blast (Altschul et al. 1997; Madeira et al. 2019), (5 iterations, 502 hits) homology searches in UniProtKB (Consortium 2018) and NCBI nr databases (Coordinators 2018)(respectively), as of March 2019. In order to extend the coverage, the searches were re-run with the lowest scoring above-the-threshold Jackhmmer hits in *Archaea* (OPX82726.1 / A0A1V4VC50_9EURY),(Nobu et al. 2017), *Chloroflexi* (ACL23627.1 / B8G4Z4_CHLAD) and *Proteobacteria* (MBN58330.1 / A0A2E7JCT6_9GAMM) (Tully et al. 2018), resulting in 2383, 2553 and 939 hits, respectively. In all cases, standard website parameters were used. Combined results consisted of 2797 non-redundant sequences including 2354 sequences shorter than 200 amino acids. The length threshold was chosen to filter out Bell homologues forming “all-in-one” NLR architectures.

### Neighboring NLRs identification

All sequence identifiers in the set were mapped to NCBI accessions using UniProt mapping tool. Accessions of identical sequences in the NCBI nr database were added using the Blast database command tool (Camacho et al. 2009). This resulted in set of 2810 protein accessions from genome-wide studies, restricted to GenBank and RefSeq NP and YP series accessions. An in-house Python (version 3.5) script (aided by requests (Reitz, K., n.d.) and xmltodict (Blech, n.d.) packages) was used to query NCBI Entrez E-utils in order to fetch almost 23k proteins coded by genes within the +/-5000 bp neighborhood of the genes encoding these Bell homologues. Among the proteins coded by the Bell-neighboring genes, 730 were matched with Pfam (El-Gebali et al. 2018), NACHT (PF05729) (Koonin and Aravind 2000) or NB-ARC (PF00931) (van der Biezen and Jones 1998) profiles. This included 467 sequences (426 unique) where the match bordered a relatively short N-terminus of 10 to 150 amino acids in length (396 unique extensions), which is a typical feature of fungal NLR proteins containing amyloid signaling motifs.

### Motif extraction

The C-terminal boundaries of the Bell domains were delimited according to the final Profile HMM (pHMM) (Eddy 2008) of the original Jackhmmer search using the hmmalign tool of the standalone HMMER distribution (version 3.2.1) (Eddy 2011). The C-terminal extensions longer than 10 amino acids were extracted. The redundancy of the set was reduced to 90% using cdhit (Fu et al. 2012; Li and Godzik 2006), (version 4.6, standard parameters) resulting in 1814 unique sequences. The common motifs in the C-termini were extracted using MEME (Bailey et al. 2009; Bailey and Elkan 1994) (version 5.0.2, standalone), allowing for motifs of any length from 6 to 30 and possibly repeated in sequence (--anr option), while requiring that each motif was found in at least 10 instances. The search yielded 66 motifs (prefixed later “all”) above the standard E-value threshold of 0.1. Aiming at motifs restricted to taxonomic branches, additional MEME searches (requiring at least 5 motif instances only) were performed for taxonomic subsets of UniProtKB sequences including: *Cyanobacteria* (found 27 motifs in 359 sequences, prefixed later “cya”), combined *Proteobacteria* and *Chloroflexi* (5 in 69, “pch”), and *Streptomycetes* (19 in 608, “str”). Finally, the MEME search with the same parameters was performed in N-termini of the NLR-containing genomic neighbors of Bell domains (39 motifs found in 338 sequences, “nlr”).

### Motif profile generation

For each motif, a pHMM was trained using hmmbuild from the HMMER package on instances reported by MEME. Then, the pHMMs were used to re-search the Bell C-terminal domains and NLR N-terminal domains, respectively, using hmmsearch with sequence and domain E-values set to 0.01 and heuristic filters turned off for sensitivity (--max option). The resulting hits were extended each side by 5 amino acids, and then re-aligned to the pHMMs and trimmed of unaligned residues using hmmalign (--trim option). Eventually, the resulting alignments were used to train final pHMMs for the motifs again using hmmbuild with standard options. The entire procedure aimed at generalizing the motifs, especially gapped, which could be truncated or over-specialized by MEME.

### Motif pairs identification

Motif pairs were identified whenever hits from the same profile HMM were found in Bell C-termini and their corresponding NLR(s) N-termini, using hmmsearch with sequence and domain E-value set to 0.01 and maximum sensitivity (--max option). There were 1157 such pairs (904 using Bell-side motifs and 253 using NLR-side motifs) in 315 (293 and 246, respectively) sequence pairs (there was a considerable overlap between motifs, see below). Only motifs with at least 5 unique pair hits (in terms of sequence) were considered. The criterion was met by 29 Bell-side motifs and 12 NLR-side motifs with 1087 hits in 295 sequence pairs.

### Motif clustering

Motifs were grouped based on overlapping matches (hits in the same sequence pairs). Motifs were joined if at least half of pairs hit by one motif profile HMM were also matched by the other. The procedure yielded ten motif classes termed BASS1 to 10. For each class, the member motif matching most sequences was used as the class representative.

### Motif characterization

For each member motif in classes BASS1-10, the sequence regions matched with its pHMM were extended each side by 5 amino acids and pairwisely locally aligned using the Waterman-Eggert method (Huang and Miller 1991; Waterman and Eggert 1987) implemented in EMBOSS (matcher tool, version 6.6.0.0) (Rice et al. 2000). The standard parameters were used including the BLOSUM62 matrix (Henikoff and Henikoff 1992) and gap opening/extending penalties 14/4. For each motif class representative, the sequence fragments from the resulting pairwise alignments were combined into multiple sequence alignments using Clustal Omega (Sievers and Higgins 2018; Sievers et al. 2011)(version 1.2.4, standalone) with the --dealign and --auto switches. The resulting MSAs were trimmed manually and profiles were generated using weblogo3 (Crooks et al. 2004) (Weblogo software repository webpage, 2019).

In addition, for each motif class and sequence, the pairwise alignments were used to obtain the combined longest fragment matching any member motif. The dealigned fragments were then combined into MSA using Clustal Omega with --full and --full-iter options, curated when necessary (BASS4) and trimmed manually. Amino acid composition of such generated MSAs was then calculated using the quantiprot package (Konopka et al. 2017), after removing redundant sequences.

The per-class and overall (unweighted) composition was then compared with amino acid composition of non-redundant sequence sets of fungal functional amyloid motifs including Het-s Related Amyloid Motif (HRAM) (Daskalov et al. 2015a), Pfam NACHT_sigma (PF17106) and Ses_B (PF17046, aka PP-motif) (Daskalov et al. 2015a; Dyrka et al. 2014), beta-solenoid repeat regions extracted from the Protein Data Bank (Berman et al. 2000) according to RepeatsDB (only reviewed entries, as of May 2019), experimentally verified amyloid fragments from AmyLoad (Wozniak and Kotulska 2015) (as of March 2017), and intrinsically disordered protein regions from DisProt (Piovesan et al. 2017) (as of July 2019, only unambiguous entries). Current SwissProt amino acid statistics were used as a reference.

### Highly internally conserved pairs search

Local pairwise alignments of C-termini of Bell domains and their neighboring NLR(s) N-termini were performed using the matcher tool with the standard parameters. For each pair ten best alternatives were filtered for minimum alignment length of 15 amino acid and minimum score of 40. The procedure yielded in 292 hits (including 234 unique) in 283 sequence pairs. The number included 44 sequence pairs not matched with the BASS1-10 motifs, comprising 25 sequence pairs not matched with pHMM of any motif.

### Phylogenetic distribution

Protein accession lists of genome assemblies listed in the Genome Taxonomy Database (GTDB) (Parks et al. 2018) metadata sheets for Bacteria (113,324 items) and Archaea (1183) were downloaded from NCBI ftp (ftp.ncbi.nlm.nih.gov/genomes/all/; as of April 2019). The accesions were matched with the ABC_tran (PF00005) (Rosteck et al. 1991), NACHT, NB-ARC, Beta_propeller (CL0186) (Murzin 1992), TPR (CL0020) (Lamb et al. 1995), CHAT (PF12770) (Aravind and Koonin 2002), PNP_UDP_1 (PF01048) (Mushegian and Koonin 1994) and TIR (PF01582) (Bonnert et al. 1997)Pfam profiles hit lists downloaded from Pfam ftp (ftp.ebi.ac.uk/pub/databases/Pfam/current_release/; as of February 2019). In addition, distributions of previously found Bell domain homologues and motif pairs involving their C-termini in GTDB-listed genomes were recored. For each, the total number of hits and the number of genomes with hits is provided. The sequence accession redundancy was resolved with the NCBI Blast nr database (as of March 2019).

### Strains and plasmids

To avoid interference with endogenous prions, the *P. anserina Δhet-s (ΔPa_3_620) Δhellf (ΔPa_3_9900) ΔPahellp (ΔPa_5_8070)* strain was used as recipient strain for the expression of molecular fusions of BASS motifs and the GFP (green fluorescent protein) or RFP (red fluorescent protein). These fusions were expressed either from a plasmid based on the pGEM-T backbone (Promega) named pOP plasmid (Daskalov *et al*. 2016), containing RFP, or from a derivative of the pAN52.1 GFP vector (Balguerie *et al*. 2004), named pGB6-GFP plasmid containing GFP. In both cases the molecular fusions were under the control of the constitutive *P. anserina* gpd (glyceraldehyde-3-phosphate dehydrogenase) promoter. The *ΔPahellp Δhet-s Δhellf* strain was transformed as described (Bergès and Barreau 1989) with one or two fusion constructs along with a vector carrying a phleomycin-resistance gene ble, pPaBle (using a 10:1 molar ratio). Phleomycin-resistant transformants were selected, grown for 30 h at 26°C and screened for the expression of the transgenes using a fluorescence microscope. Fragments (protein position indicated in brackets) of the following genes (accessions from GenBank or RefSeq) were amplified using specific primers: CAB66307.1 (110-139), CAB66306.1 (1-34), WP_037701008.1 (70-124), WP_037701012.1 (1-37), ACC79696.1 (94-126), ACC79697.1 (1-38), WP_063130184.1 (74-128). The PCR products were cloned upstream of the RFP coding sequence in the pOP plasmid using *Pac*I/*Bgl*II restriction enzymes or downstream of the GFP in the pAN52.1 plasmid using *Not*I/*Bam*HI restriction enzymes. For heterologous expression in *E. coli*, the following fragments were amplified using specific primers and cloned in pET24a (Novagen) using the *Nde*I/*Xho*I restrictions sites: CAB66307.1 (110-139) or (38-139), WP_037701008.1 (97-124), ACC79696.1 (94-126), WP_063130184.1 (100-128).

Since the sequences selected for the BASS3 motif were not included neither in the GenBank nor RefSeq NP or YP series at the time of the genome mining, they were not processed for identifying neighboring NLRs, and hence are not covered in the motif pair list in Table S2. Nevertheless, the BASS3 motifs can be found in these sequences with the motif pHMMs at default identification threshold (E-value of 0.01), except for ACC79697.1 being slightly below (E-value of 0.1).

### Prion propagation

The [b] phenotype (acquisition of the [b] prion) was defined in *Podospora* strains expressing fluorescent fusion proteins as the absence ([b*]) or the presence ([b]) of fluorescent dot-like aggregates. To monitor the propagation of the [b] prion, prion free strains were subcultured in presence of [b] prion donor strain and after 72h (contact between tested and donor strains was established after 24h of subculture) the initially prion free tested strain was subcultured on fresh medium and monitored for the presence of aggregates using fluorescence microscopy.

### Protein preparation and fibril formation

6his-tagged proteins were expressed from pET24a constructs in *E. coli* BL21-CodonPlus®-RP competent cells as insoluble proteins and purified under denaturing conditions using its terminal 6 histidine tag as previously described (Dos Reis *et al*. 2002). Briefly, cells were grown at 37°C in DYT medium to 0.6 OD_600_ and expression was induced with 1 mM isopropyl β-D-1-thiogalactopyranoside. After, 4 h, cells were harvested by centrifugation, frozen at - 80°C sonicated on ice in a lysis buffer (Tris 50 mM, 150 mM NaCl, pH 8) and centrifuged for 20 min at 20,000 g to remove *E. coli* contaminants. The pellet was washed in the same buffer and resuspended in denaturing buffer (6M guanidinium HCl, 150 mM NaCl, and 100 mM Tris-HCl, pH 8) until complete solubilization. The lysate was incubated with Talon Resin (CLONTECH) for 1 h at 20°C, and the resin was extensively washed with 8 M urea, 150 mM NaCl, and 100 mM Tris-HCl, pH 8. The protein was eluted from the resin in the same buffer containing 200 mM imidazole. The proteins were pure as judged by sodium-dodecyl-sulfate polyacrylamide-gel electrophoreses (SDS-PAGE) followed by Coomassie-Blue staining and yield was in the range of ∼2-4 mg of protein per liter of culture. To eliminate urea, elution buffer was replaced by overnight dialysis at 4°C against Milli-Q water. Fibrills formation resulted spontaneously from dialysis process followed by sample storage in H_2_0 or in ammonium acetate buffer 100 mM pH 4.5 at 4°C for 7 to 30 days.

### Light Microscopy

*P. anserina* hyphae were inoculated on solid medium and cultivated for 24 to 48 h at 26°C. The medium was then cut out, placed on a glass slide and examined with a Leica DMRXA microscope equipped with a Micromax CCD (Princeton Instruments) controlled by the Metamorph 5.06 software (Roper Scientific). The microscope was fitted with a Leica PL APO 100X immersion lens.

### Transmission Electron Microscopy

For fibrils observations, negative staining was performed: aggregated proteins were adsorbed onto Formvar-coated copper grids (400 mesh) and allowed to dry for 15 min in air, grids were then negatively stained 1 min with 10 μL of freshly prepared 2% uranyl acetate in water, dried with filter paper, and examined with a Hitachi H7650 transmission electron microscope (Hitachi, Krefeld, Germany) at an accelerating voltage of 120 kV. TEM was performed at the Pôle Imagerie Électronique of the Bordeaux Imaging Center using a Gatan USC1000 2k x 2k camera.

### Solid-state NMR of BASS1 fibrils

The solid-state NMR spectrum of BASS 1 was recorded at 800 MHz on a Bruker Biospin magnet using a triple resonance 3.2 mm probe at a spinning frequency of 11 kHz. 64 scans were used using a recycle delay of 3 sec and an acquisition time of 17 ms.

### X-ray diffraction of BASS1 fibrils

Fiber diffraction pattern was measured at 4°C on an Excillum MetalJet X-ray generator at the galium wavelength (Kα, λ = 1.34 Å). The source was equipped with Xenocs Fox3D optics and a Dectris Eiger 1M detector on a 2θ arm of a STOE stadivari goniometer. The detector has been turned by 90° to have the blind region vertical to hide as much as possible the shadow of the beamstop. The viscous concentrated hydrated sample was mounted in a MicroLoop™ from Mitegen on a goniometer head under the cold nitrogen flow. The diffraction pattern corresponds to a 360° rotation along the phi axis with an exposure time of 180 sec.

---

**Figure S1.**
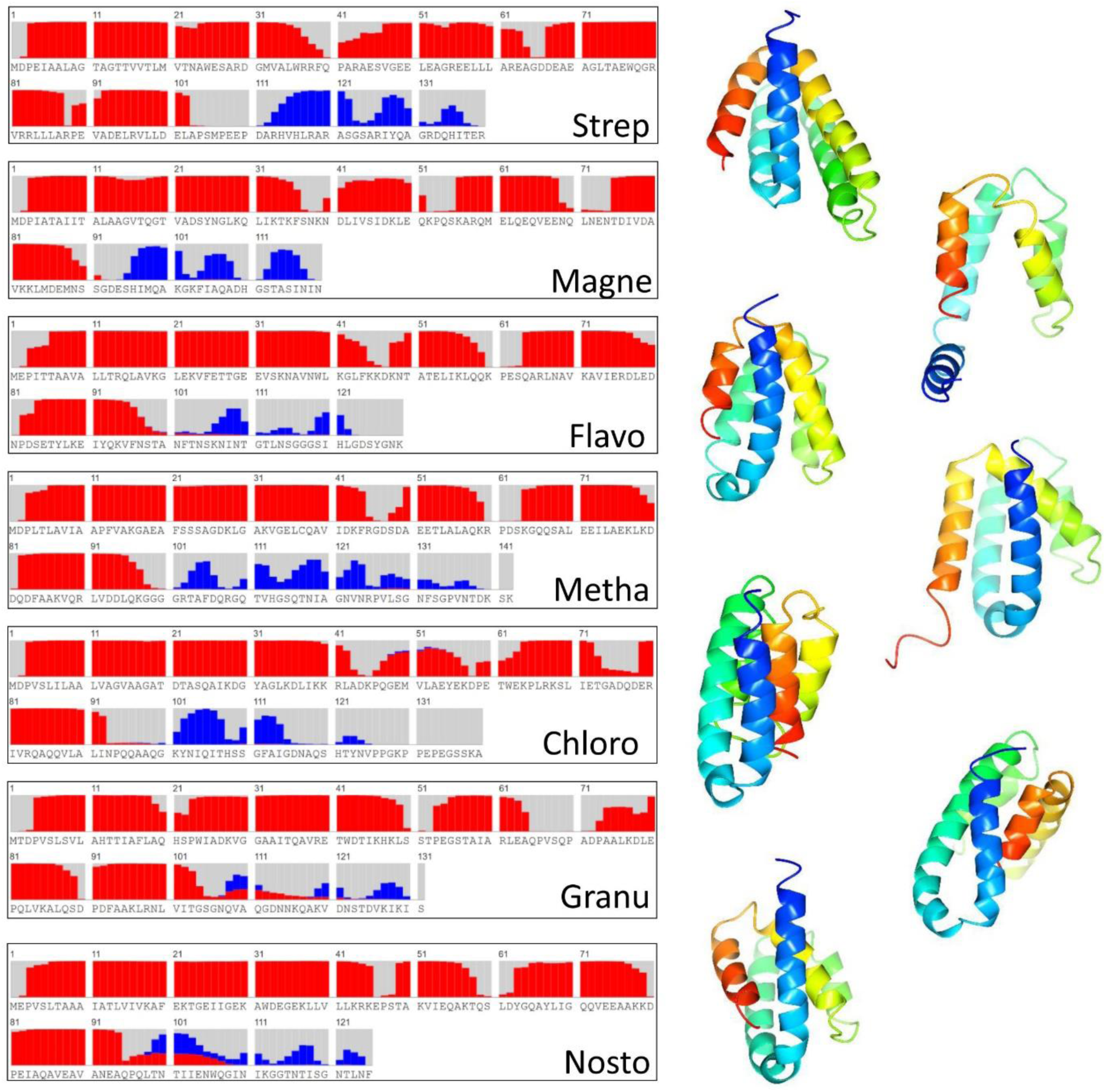
Homology models of Bell-domain proteins. Secondary structure prediction and homology models are given for a set of phylogenetically diverse Bell-domain proteins from prokaryotes ((Strep, Q9RDG0 from Streptomyces coelicolor A3(2); Chlor, HBY96210.1 from *Chloroflexi bacterium*; Sacch, ONI86675.1 from *Saccharothrix sp. ALI-22-I*; Flavo, SDZ50707.1 from *Flavobacterium aquidurense*; Nosto, RCJ33357.1 from *Nostoc punctiforme NIES-2108*; Granu, ROP69996.1 from *Granulicella sp. GAS466*; Metha, AEB69174.1 from *Methanothrix soehngenii (strain ATCC 5969)*; Magne, ETR68090.1 from *Candidatus Magnetoglobus multicellularis str. Araruama*). Secondary structure prediction and homology models are as given by RAPTOR-X contact. Red bars represent α-helical propensity, blue bar β-sheet propensity. Secondary structure prediction are given for the full-length protein, the homology model for the Bell-domain region only. Note that secondary structure prediction often merged helix 1 and 2 which in some cases are modelled as a continuous helix.

**Figure S2.**
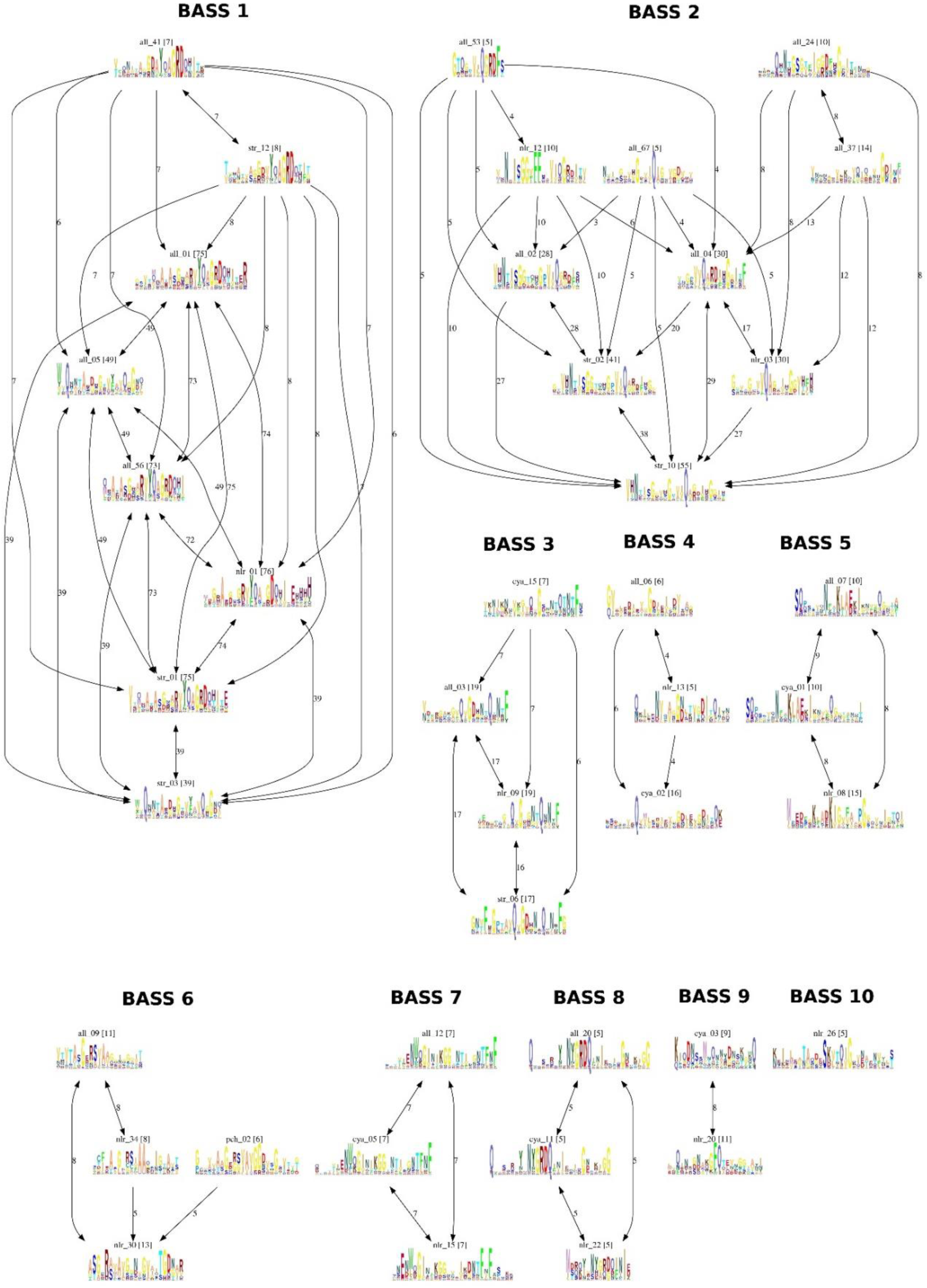
Clustering of motifs. Clustering of motifs. 29 motifs matching at least 5 non-redundant pairs of NLRs and Bell-domain proteins were clustered based on overlapping matches. Motifs (presented as profile HMM logos) were joined if at least half of the pairs hit by one motif profile HMM were also matched by the other. The number in brackets represents the number of pairs for each profile, the number on the arrow joining the motifs indicates the number of common matching underlying sequences. The grouping was preserved in an alternative clustering scheme where motifs sharing at least 40% of their underlying sequences were grouped together.

**Figure S3.**
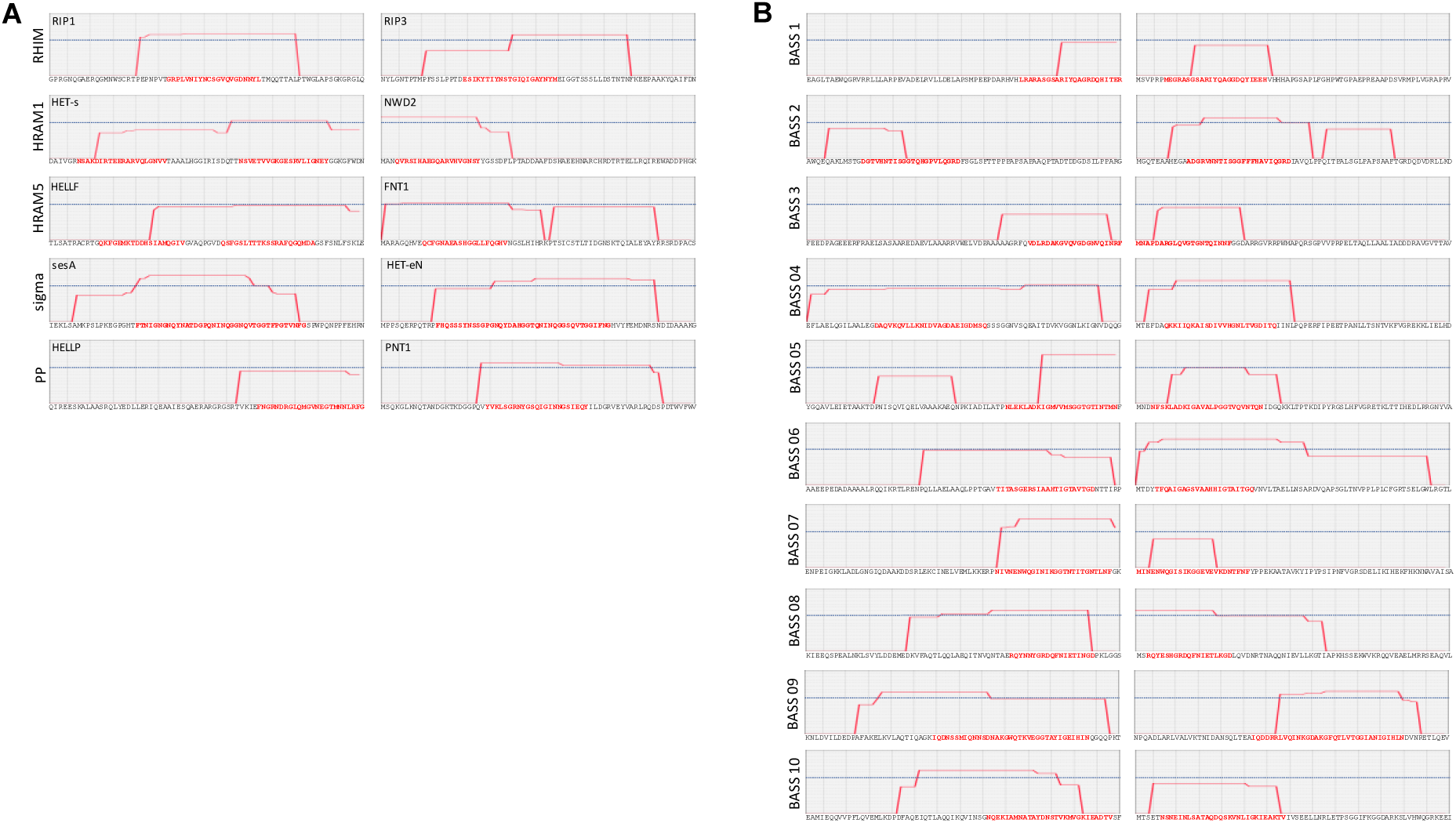
Amyloid formation propensity of the different motifs based on ArchCandy. **A.** ArchCandy amyloid propensity for various amyloid signaling motifs from mammals and fungi are given. Position of the amyloid motif is highlighted in red within a 70 amino acid long sequence window. The ArchCandy prediction is given in red, the blue line gives the recommend significance threshold. For the fungal motifs, prediction of amyloid propensity in the effector domain-associated motif is given in the left column and that of the NLR-associated motif encoded by the adjacent gene is given in the right column**. B.** ArchCandy amyloid propensity is representative protein pairs corresponding to the 10 BASS motifs are give; predictions for the Bell-domain associated motif (left) and the NLR-associated motif (right) are given in the same way as in A. Gene numbers and species names of the representative pairs are as given in Fig. 2.

**Figure S4.**
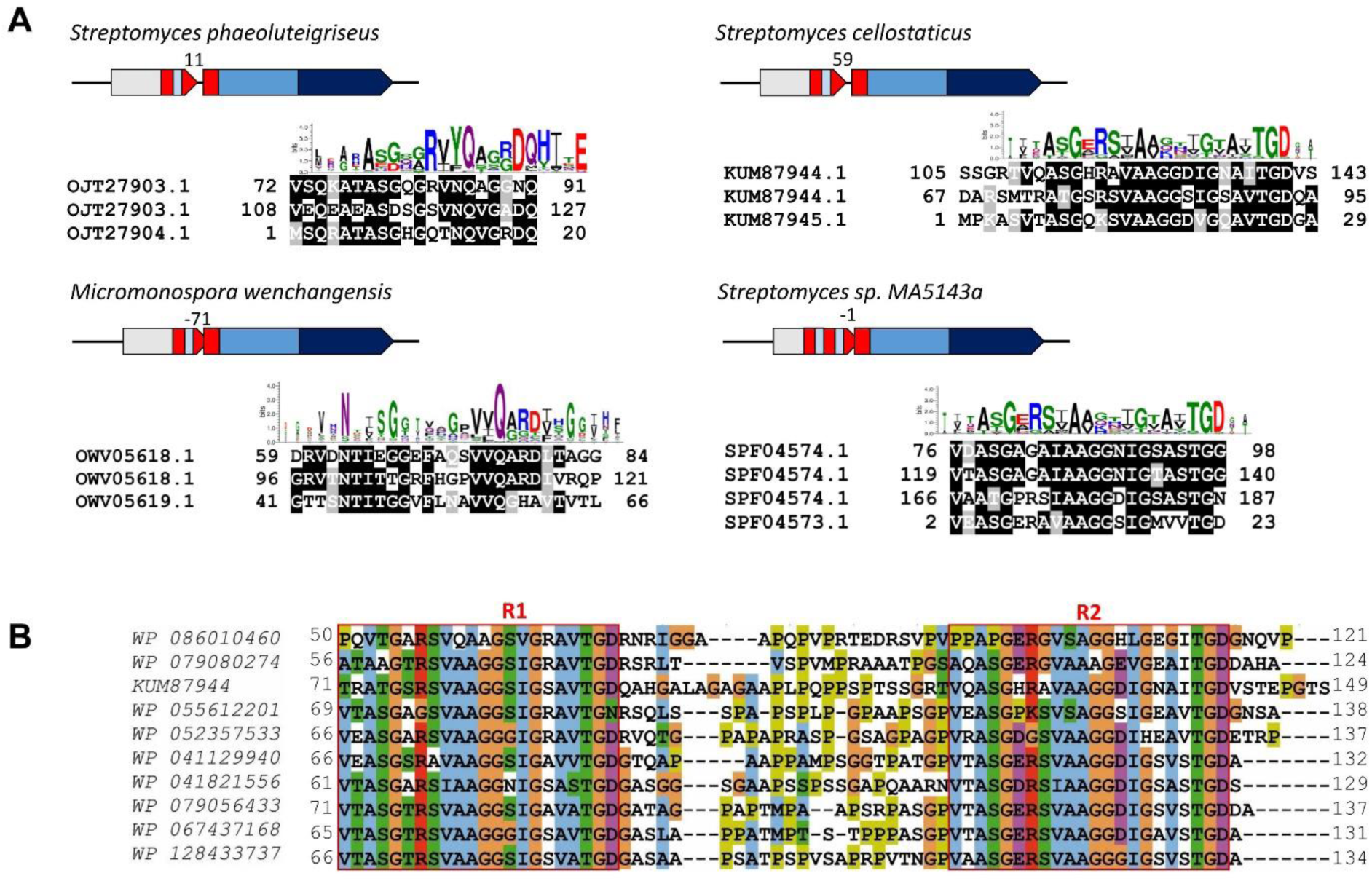
Double and triple BASS motifs. **A.** Species of origin, gene identification and gene architecture of selected pairs of gene encoding a Bell domain and a NLR and sharing an amyloid signaling motif are given together with an alignment of the two (or three motifs) found associated to the Bell domain with the motif found associated to the NLR. The consensus signature sequence of the motif is given above the alignment. The sequences encoding the BASS motifs are represented in red, the Bell-domain in grey, NB-ARC domain is light blue, TPR repeats in dark blue. **B.** Alignment of 10 orthologs of KUM87944.1 protein from *Streptomyces cellostaticus* given in A, showing the two repeats of the motif and the variable proline and glycine-rich region between the R1 and R2 repeats.

**Figure S5.**
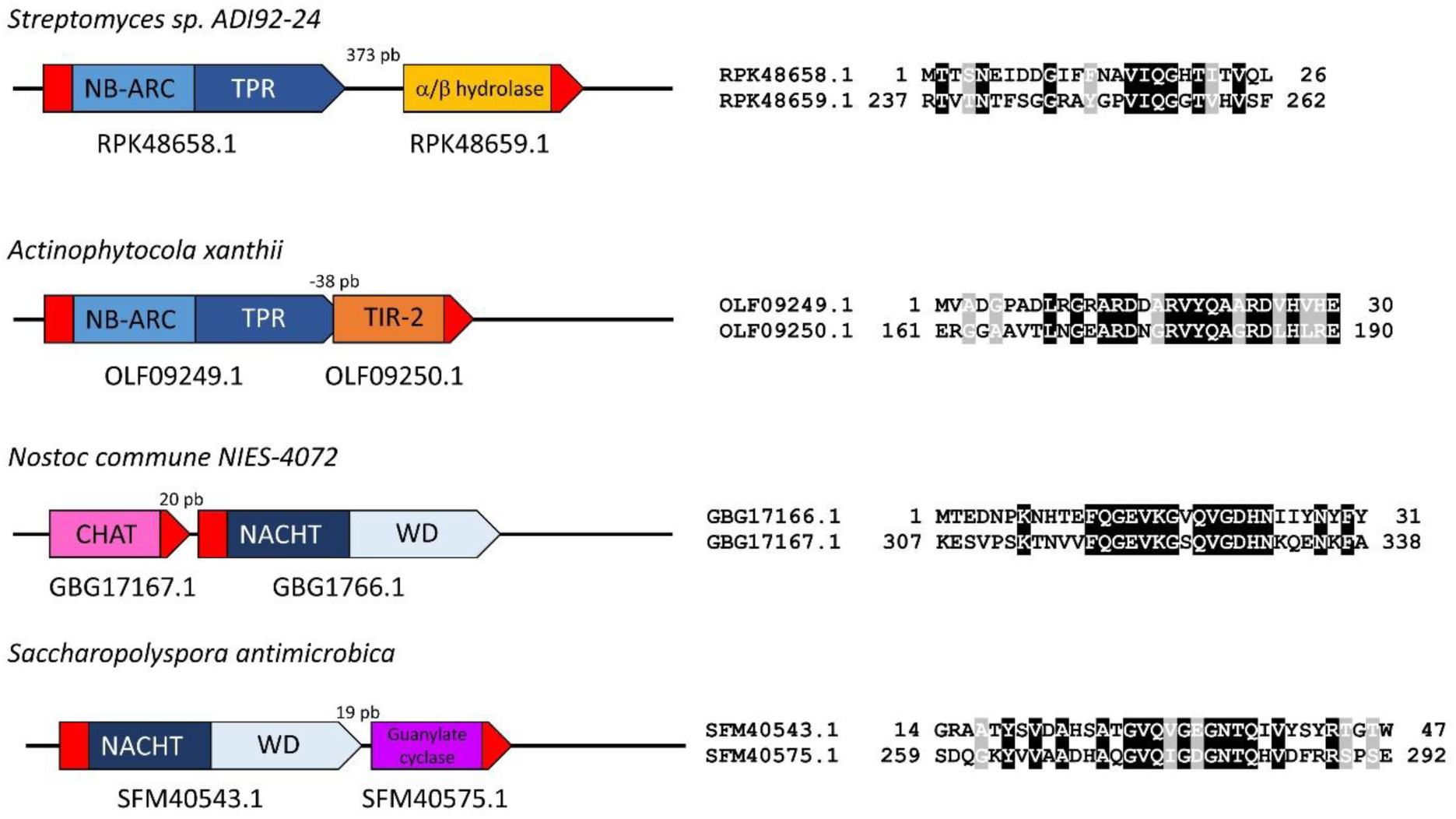
Other BASS-associated effector domains. Examples of gene pairs sharing an amyloid signaling motif in which the effector domain is not a Bell domain. In each case, the species of origin, gene identification and gene architecture of the selected pairs is given together with an alignment of the motifs associated to the effector domain and associated to the NLR. The sequences encoding the BASS motif are represented in red.

**Figure S6.**
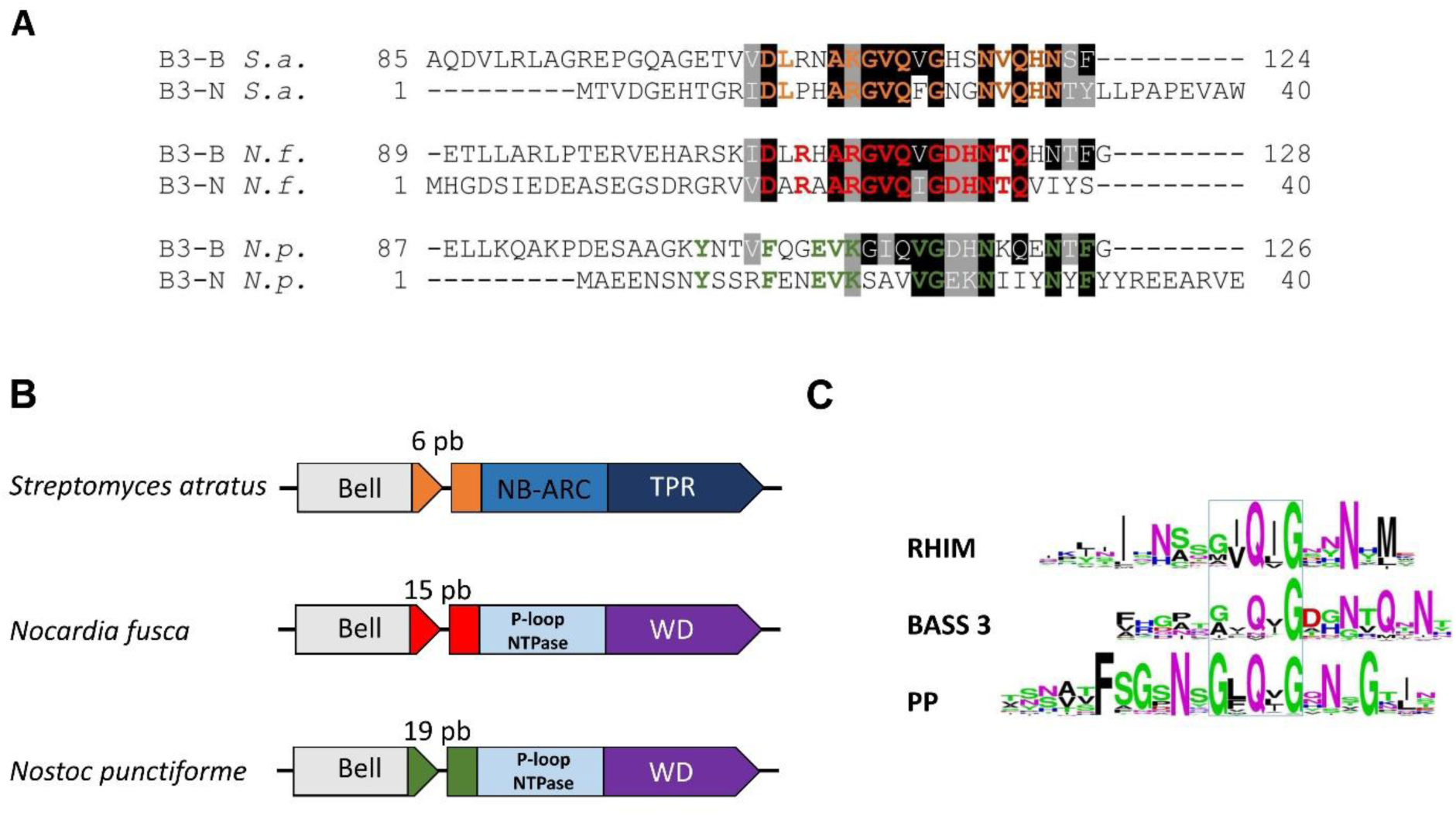
BASS3 motif sequences selected for *in vivo* and *in vitro* expression studies. **A.** Alignement of the BASS3 motif of the selected proteins, the grey and black boxing given residues similar or identical respectively in at least 4 of the 6 sequences. The colored residues highlight identical residues within a gene pair, that is, identical in the Bell (B) and NLR-side (N) motif. Accession number of the sequences are given Table 3. **B.** Species of origin and genome architecture are given for the three selected the Bell-domain and NLR pairs. The number given above the gene diagram is the distance between the Bell-domain encoding and NLR encoding ORF. C. Comparaison of the consensus sequences of the metazoan RHIM, bacterial BASS 3 and fungal PP-motifs. Logos where generated using all RHIM Pfam entries (PF17721) from Metazoans, all identified BASS 3 pairs motifs (Table S2) and all Pfam entries for PP (PF17046).

**Figure S7.**
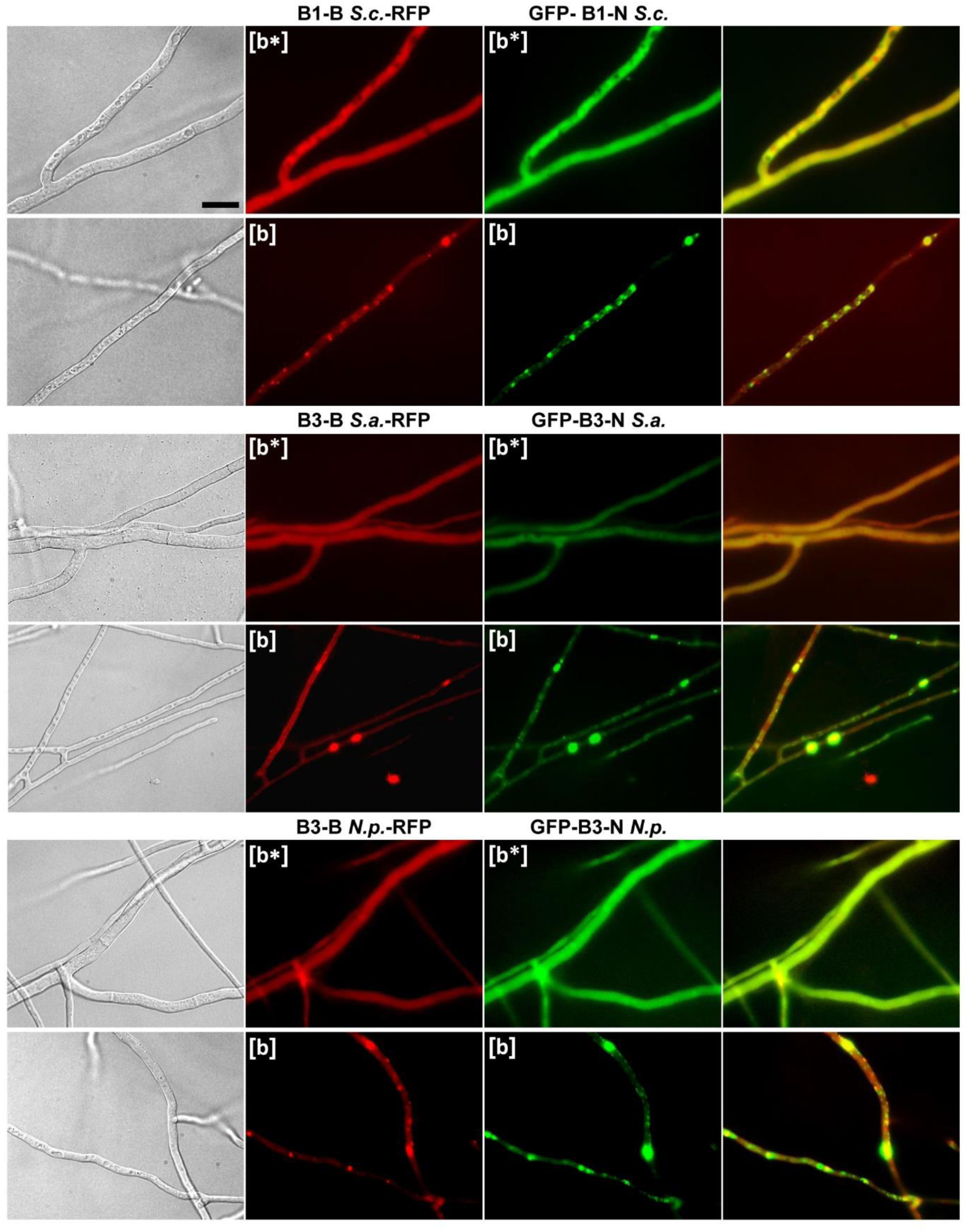
Bell and NLR-side BASS motif co-aggregation in *Podospora anserina*. Micrographs of *Podospora anserina* strains co-expressing Bell-side BASS motifs fused to RFP (in C-terminus) as indicated and the corresponding NLR-side motif fused to GFP (in N-terminus), (Scale bar 5 μm). Panel are from left to right, bright field, RFP, GFP and overlay.

## References

Adachi H, Contreras M, Harant A, Wu CH, Derevnina L, Sakai T, Duggan C, Moratto E, Bozkurt TO, Maqbool A, et al. (2019). An N-terminal motif in NLR immune receptors is functionally conserved across distantly related plant species. eLife 8.

Ahmed AB, and Kajava AV (2013). Breaking the amyloidogenicity code: Methods to predict amyloids from amino acid sequence. Febs Letters 587, 1089–1095.

Ahmed AB, Znassi N, Chateau MT, and Kajava AV (2014). A structure-based approach to predict predisposition to amyloidosis. Alzheimer’s & dementia : the journal of the Alzheimer’s Association.

Altschul SF, Madden TL, Schäffer AA, Zhang J, Zhang Z, Miller W, and Lipman DJ (1997). Gapped BLAST and PSI-BLAST: a new generation of protein database search programs. Nucleic Acids Research 25, 3389–3402.

Aravind L, and Koonin EV (2002). Classification of the caspase-hemoglobinase fold: detection of new families and implications for the origin of the eukaryotic separins. Proteins 46, 355–367.

Asplund-Samuelsson J, Bergman B, and Larsson J (2012). Prokaryotic caspase homologs: phylogenetic patterns and functional characteristics reveal considerable diversity. PLoS One 7, e49888.

Bailey TL, Boden M, Buske FA, Frith M, Grant CE, Clementi L, Ren J, Li WW, and Noble WS (2009). MEME Suite: tools for motif discovery and searching. Nucleic Acids Research 37, W202–W208.

Bailey TL, and Elkan C (1994). Fitting a mixture model by expectation maximization to discover motifs in biopolymers. Proceedings International Conference on Intelligent Systems for Molecular Biology 2, 28–36.

Balguerie A, Dos Reis S, Ritter C, Chaignepain S, Coulary-Salin B, Forge V, Bathany K, Lascu I, Schmitter JM, Riek R, et al. (2003). Domain organization and structure-function relationship of the HET-s prion protein of Podospora anserina. Embo J 22, 2071–2081.

Baxa U, Cassese T, Kajava AV, and Steven AC (2006). Structure, function, and amyloidogenesis of fungal prions: filament polymorphism and prion variants. Adv Protein Chem 73, 125–180.

Benkemoun L, Ness F, Sabate R, Ceschin J, Breton A, Clave C, and Saupe SJ (2011). Two structurally similar fungal prions efficiently cross-seed in vivo but form distinct polymers when coexpressed. Mol Microbiol 82, 1392–1405.

Benkemoun L, Sabate R, Malato L, Dos Reis S, Dalstra H, Saupe SJ, and Maddelein ML (2006). Methods for the in vivo and in vitro analysis of [Het-s] prion infectivity. Methods 39, 61–67.

Berman HM, Westbrook J, Feng Z, Gilliland G, Bhat TN, Weissig H, Shindyalov IN, and Bourne PE (2000). The Protein Data Bank. Nucleic Acids Research 28, 235–242.

Bonnert TP, Garka KE, Parnet P, Sonoda G, Testa JR, and Sims JE (1997). The cloning and characterization of human MyD88: a member of an IL-1 receptor related family. FEBS letters 402, 81–84.

Cai X, Chen J, Xu H, Liu S, Jiang QX, Halfmann R, and Chen ZJ (2014). Prion-like polymerization underlies signal transduction in antiviral immune defense and inflammasome activation. Cell 156, 1207–1222.

Cai X, Xu H, and Chen ZJ (2016). Prion-Like Polymerization in Immunity and Inflammation. Cold Spring Harb Perspect Biol.

Camacho C, Coulouris G, Avagyan V, Ma N, Papadopoulos J, Bealer K, and Madden TL (2009). BLAST+: architecture and applications. BMC bioinformatics 10, 421.

Chevanne D, Bastiaans E, Debets A, Saupe SJ, Clave C, and Paoletti M (2009). Identification of the het-r vegetative incompatibility gene of Podospora anserina as a member of the fast evolving HNWD gene family. Curr Genet 55, 93–102.

Chevanne D, Saupe SJ, Clave C, and Paoletti M (2010). WD-repeat instability and diversification of the Podospora anserina hnwd non-self recognition gene family. BMC Evol Biol 10, 134.

Choi GH, Dawe AL, Churbanov A, Smith ML, Milgroom MG, and Nuss DL (2012). Molecular characterization of vegetative incompatibility genes that restrict hypovirus transmission in the chestnut blight fungus Cryphonectria parasitica. Genetics 190, 113–127.

Colby DW, and Prusiner SB (2011). Prions. Cold Spring Harb Perspect Biol 3, a006833.

Consortium TU (2018). UniProt: a worldwide hub of protein knowledge. Nucleic Acids Research 47, D506–D515.

Coordinators NR (2018). Database resources of the National Center for Biotechnology Information. Nucleic Acids Research 46, D8–D13.

Coustou-Linares V, Maddelein ML, Begueret J, and Saupe SJ (2001). In vivo aggregation of the HET-s prion protein of the fungus Podospora anserina. Mol Microbiol 42, 1325–1335.

Crooks GE, Hon G, Chandonia J-M, and Brenner SE (2004). WebLogo: a sequence logo generator. Genome Research 14, 1188–1190.

Daskalov A, Dyrka W, and Saupe SJ (2015a). Theme and variations: evolutionary diversification of the HET-s functional amyloid motif. Scientific reports 5, 12494.

Daskalov A, Habenstein B, Martinez D, Debets AJ, Sabate R, Loquet A, and Saupe SJ (2015b). Signal transduction by a fungal NOD-like receptor based on propagation of a prion amyloid fold. PLoS Biol 13, e1002059.

Daskalov A, Habenstein B, Sabate R, Berbon M, Martinez D, Chaignepain S, Coulary-Salin B, Hofmann K, Loquet A, and Saupe SJ (2016). Identification of a novel cell death-inducing domain reveals that fungal amyloid-controlled programmed cell death is related to necroptosis. Proc Natl Acad Sci U S A 113, 2720–2725.

Daskalov A, Paoletti M, Ness F, and Saupe SJ (2012). Genomic clustering and homology between HET-S and the NWD2 STAND protein in various fungal genomes. Plos One 7, e34854.

Durand PM, Sym S, and Michod RE (2016). Programmed Cell Death and Complexity in Microbial Systems. Curr Biol 26, R587–R593.

Dyrka W, Lamacchia M, Durrens P, Kobe B, Daskalov A, Paoletti M, Sherman DJ, and Saupe SJ (2014). Diversity and Variability of NOD-Like Receptors in Fungi. Genome biology and evolution 6, 3137–3158.

Eddy SR (2008). A probabilistic model of local sequence alignment that simplifies statistical significance estimation. PLoS computational biology 4, e1000069.

Eddy SR (2011). Accelerated Profile HMM Searches. PLoS computational biology 7, e1002195.

El-Gebali S, Mistry J, Bateman A, Eddy SR, Luciani A, Potter SC, Qureshi M, Richardson LJ, Salazar GA, Smart A, et al. (2018). The Pfam protein families database in 2019. Nucleic Acids Research 47, D427–D432.

Erskine E, MacPhee CE, and Stanley-Wall NR (2018). Functional Amyloid and Other Protein Fibers in the Biofilm Matrix. J Mol Biol 430, 3642–3656.

Espagne E, Balhadere P, Penin ML, Barreau C, and Turcq B (2002). HET-E and HET-D Belong to a New Subfamily of WD40 Proteins Involved in Vegetative Incompatibility Specificity in the Fungus Podospora anserina. Genetics 161, 71–81.

Freihat LA, Wheeler JI, Wong A, Turek I, Manallack DT, and Irving HR (2019). IRAK3 modulates downstream innate immune signalling through its guanylate cyclase activity. Scientific reports 9, 15468.

Fu L, Niu B, Zhu Z, Wu S, and Li W (2012). CD-HIT: accelerated for clustering the next-generation sequencing data. Bioinformatics (Oxford, England) 28, 3150–3152.

Fukuda Y, Nakayama Y, and Tomita M (2003). On dynamics of overlapping genes in bacterial genomes. Gene 323, 181–187.

Giraldo R, Moreno-Diaz de la Espina S, Fernandez-Tresguerres ME, and Gasset-Rosa F (2011). RepA-WH1 prionoid: a synthetic amyloid proteinopathy in a minimalist host. Prion 5, 60–64.

Graziani S, Silar P, and Daboussi MJ (2004). Bistability and hysteresis of the ‘Secteur’ differentiation are controlled by a two-gene locus in Nectria haematococca. BMC Biol 2, 18.

Greenwald J, Buhtz C, Ritter C, Kwiatkowski W, Choe S, Maddelein ML, Ness F, Cescau S, Soragni A, Leitz D, et al. (2010). The mechanism of prion inhibition by HET-S. Mol Cell 38, 889–899.

Heller J, Clave C, Gladieux P, Saupe SJ, and Glass NL (2018). NLR surveillance of essential SEC-9 SNARE proteins induces programmed cell death upon allorecognition in filamentous fungi. Proc Natl Acad Sci U S A.

Henikoff S, and Henikoff JG (1992). Amino acid substitution matrices from protein blocks. Proceedings of the National Academy of Sciences 89, 10915–10919.

Hu XJ, Li T, Wang Y, Xiong Y, Wu XH, Zhang DL, Ye ZQ, and Wu YD (2017). Prokaryotic and Highly-Repetitive WD40 Proteins: A Systematic Study. Scientific reports 7, 10585.

Huang X, and Miller W (1991). A Time-Efficient, Linear-Space Local Similarity Algorithm. Advances in Applied Mathematics 12, 337–357.

Hug LA, Baker BJ, Anantharaman K, Brown CT, Probst AJ, Castelle CJ, Butterfield CN, Hernsdorf AW, Amano Y, Ise K, et al. (2016). A new view of the tree of life. Nat Microbiol 1, 16048.

Iranzo J, Lobkovsky AE, Wolf YI, and Koonin EV (2014). Virus-host arms race at the joint origin of multicellularity and programmed cell death. Cell Cycle 13, 3083–3088.

Jones JD, Vance RE, and Dangl JL (2016). Intracellular innate immune surveillance devices in plants and animals. Science 354.

Kajava AV, Klopffleisch K, Chen S, and Hofmann K (2014). Evolutionary link between metazoan RHIM motif and prion-forming domain of fungal heterokaryon incompatibility factor HET-s/HET-s. Scientific reports 4, 7436.

Keller NP (2019). Fungal secondary metabolism: regulation, function and drug discovery. Nat Rev Microbiol 17, 167–180.

Kleino A, Ramia NF, Bozkurt G, Shen Y, Nailwal H, Huang J, Napetschnig J, Gangloff M, Chan FK, Wu H, et al. (2017). Peptidoglycan-Sensing Receptors Trigger the Formation of Functional Amyloids of the Adaptor Protein Imd to Initiate Drosophila NF-kappaB Signaling. Immunity 47, 635–647 e636.

Konopka BM, Marciniak M, and Dyrka W (2017). Quantiprot - a Python package for quantitative analysis of protein sequences. BMC bioinformatics 18, 339.

Koonin EV, and Aravind L (2000). The NACHT family - a new group of predicted NTPases implicated in apoptosis and MHC transcription activation. Trends in Biochemical Sciences 25, 223–224.

Koonin EV, and Aravind L (2002). Origin and evolution of eukaryotic apoptosis: the bacterial connection. Cell Death Differ 9, 394–404.

Koonin EV, and Krupovic M (2019). Origin of programmed cell death from antiviral defense? Proc Natl Acad Sci U S A 116, 16167–16169.

Lamb JR, Tugendreich S, and Hieter P (1995). Tetratrico peptide repeat interactions: to TPR or not to TPR? Trends in Biochemical Sciences 20, 257–259.

Li J, McQuade T, Siemer AB, Napetschnig J, Moriwaki K, Hsiao YS, Damko E, Moquin D, Walz T, McDermott A, et al. (2012). The RIP1/RIP3 necrosome forms a functional amyloid signaling complex required for programmed necrosis. Cell 150, 339–350.

Li W, and Godzik A (2006). Cd-hit: a fast program for clustering and comparing large sets of protein or nucleotide sequences. Bioinformatics (Oxford, England) 22, 1658–1659.

Loquet A, El Mammeri N, Stanek J, Berbon M, Bardiaux B, Pintacuda G, and Habenstein B (2018a). 3D structure determination of amyloid fibrils using solid-state NMR spectroscopy. Methods 138–139, 26-38.

Loquet A, and Saupe SJ (2017). Diversity of Amyloid Motifs in NLR Signaling in Fungi. Biomolecules 7.

Loquet A, Saupe SJ, and Romero D (2018b). Functional Amyloids in Health and Disease. J Mol Biol 430, 3629–3630.

Madeira F, Park Ym, Lee J, Buso N, Gur T, Madhusoodanan N, Basutkar P, Tivey ARN, Potter SC, Finn RD, et al. (2019). The EMBL-EBI search and sequence analysis tools APIs in 2019. Nucleic Acids Research 47, W636–W641.

Marold JD, Kavran JM, Bowman GD, and Barrick D (2015). A Naturally Occurring Repeat Protein with High Internal Sequence Identity Defines a New Class of TPR-like Proteins. Structure 23, 2055–2065.

Mermigka G, Amprazi M, Mentzelopoulou A, Amartolou A, and Sarris PF (2019). Plant and Animal Innate Immunity Complexes: Fighting Different Enemies with Similar Weapons. Trends Plant Sci.

Mompean M, Li W, Li J, Laage S, Siemer AB, Bozkurt G, Wu H, and McDermott AE (2018). The Structure of the Necrosome RIPK1-RIPK3 Core, a Human Hetero-Amyloid Signaling Complex. Cell 173, 1244–1253 e1210.

Murphy JM, Czabotar PE, Hildebrand JM, Lucet IS, Zhang JG, Alvarez-Diaz S, Lewis R, Lalaoui N, Metcalf D, Webb AI, et al. (2013). The pseudokinase MLKL mediates necroptosis via a molecular switch mechanism. Immunity 39, 443–453.

Murzin AG (1992). Structural principles for the propeller assembly of beta-sheets: the preference for seven-fold symmetry. Proteins 14, 191–201.

Mushegian AR, and Koonin EV (1994). Unexpected sequence similarity between nucleosidases and phosphoribosyltransferases of different specificity. Protein Science: A Publication of the Protein Society 3, 1081–1088.

Nimma S, Ve T, Williams SJ, and Kobe B (2017). Towards the structure of the TIR-domain signalosome. Curr Opin Struct Biol 43, 122–130.

Nobu MK, Narihiro T, Liu M, Kuroda K, Mei R, and Liu W-T (2017). Thermodynamically diverse syntrophic aromatic compound catabolism. Environmental Microbiology 19, 4576–4586.

O’Neill LA, and Bowie AG (2007). The family of five: TIR-domain-containing adaptors in Toll-like receptor signalling. Nat Rev Immunol 7, 353–364.

Otzen D, and Riek R (2019). Functional Amyloids. Cold Spring Harb Perspect Biol.

Paoletti M, Saupe SJ, and Clave C (2007). Genesis of a fungal non-self recognition repertoire. Plos One 2, e283.

Parks DH, Chuvochina M, Waite DW, Rinke C, Skarshewski A, Chaumeil PA, and Hugenholtz P (2018). A standardized bacterial taxonomy based on genome phylogeny substantially revises the tree of life. Nat Biotechnol 36, 996–1004.

Pham CL, Shanmugam N, Strange M, O’Carroll A, Brown JW, Sierecki E, Gambin Y, Steain M, and Sunde M (2019). Viral M45 and necroptosis-associated proteins form heteromeric amyloid assemblies. EMBO Rep 20.

Piovesan D, Tabaro F, Mičetić I, Necci M, Quaglia F, Oldfield CJ, Aspromonte MC, Davey NE, Davidović R, Dosztányi Z, et al. (2017). DisProt 7.0: a major update of the database of disordered proteins. Nucleic Acids Research 45, D219–D227.

Potter SC, Luciani A, Eddy SR, Park Y, Lopez R, and Finn RD (2018). HMMER web server: 2018 update. Nucleic Acids Research 46, W200–W204.

Rebsamen M, Heinz LX, Meylan E, Michallet MC, Schroder K, Hofmann K, Vazquez J, Benedict CA, and Tschopp J (2009). DAI/ZBP1 recruits RIP1 and RIP3 through RIP homotypic interaction motifs to activate NF-kappaB. EMBO Rep 10, 916–922.

Rice P, Longden I, and Bleasby A (2000). EMBOSS: The European Molecular Biology Open Software Suite. Trends in Genetics 16, 276–277.

Riek R, and Eisenberg DS (2016). The activities of amyloids from a structural perspective. Nature 539, 227–235.

Ritter C, Maddelein ML, Siemer AB, Luhrs T, Ernst M, Meier BH, Saupe SJ, and Riek R (2005). Correlation of structural elements and infectivity of the HET-s prion. Nature 435, 844–848.

Rosteck PR, Reynolds PA, and Hershberger CL (1991). Homology between proteins controlling Streptomyces fradiae tylosin resistance and ATP-binding transport. Gene 102, 27–32.

Rouse SL, Matthews SJ, and Dueholm MS (2018). Ecology and Biogenesis of Functional Amyloids in Pseudomonas. J Mol Biol 430, 3685–3695.

Sabate R, Baxa U, Benkemoun L, Sanchez de Groot N, Coulary-Salin B, Maddelein ML, Malato L, Ventura S, Steven AC, and Saupe SJ (2007). Prion and non-prion amyloids of the HET-s prion forming domain. J Mol Biol 370, 768–783.

Saupe S, Turcq B, and Begueret J (1995). A gene responsible for vegetative incompatibility in the fungus Podospora anserina encodes a protein with a GTP-binding motif and G beta homologous domain. Gene 162, 135–139.

Seuring C, Greenwald J, Wasmer C, Wepf R, Saupe SJ, Meier BH, and Riek R (2012). The mechanism of toxicity in HET-S/HET-s prion incompatibility. PLoS Biol 10, e1001451.

Shahnawaz M, Park KW, Mukherjee A, Diaz-Espinoza R, and Soto C (2017). Prion-like characteristics of the bacterial protein Microcin E492. Scientific reports 7, 45720.

Shih PM, Wu D, Latifi A, Axen SD, Fewer DP, Talla E, Calteau A, Cai F, Tandeau de Marsac N, Rippka R, et al. (2013). Improving the coverage of the cyanobacterial phylum using diversity-driven genome sequencing. Proc Natl Acad Sci U S A 110, 1053–1058.

Siemer AB, Ritter C, Ernst M, Riek R, and Meier BH (2005). High-resolution solid-state NMR spectroscopy of the prion protein HET-s in its amyloid conformation. Angew Chem Int Ed Engl 44, 2441–2444.

Sievers F, and Higgins DG (2018). Clustal Omega for making accurate alignments of many protein sequences. Protein Science 27, 135–145.

Sievers F, Wilm A, Dineen D, Gibson TJ, Karplus K, Li W, Lopez R, McWilliam H, Remmert M, Söding J, et al. (2011). Fast, scalable generation of high-quality protein multiple sequence alignments using Clustal Omega. Molecular systems biology 7, 539.

Sun X, Yin J, Starovasnik MA, Fairbrother WJ, and Dixit VM (2002). Identification of a novel homotypic interaction motif required for the phosphorylation of receptor-interacting protein (RIP) by RIP3. J Biol Chem 277, 9505–9511.

Sunde M, Serpell LC, Bartlam M, Fraser PE, Pepys MB, and Blake CC (1997). Common core structure of amyloid fibrils by synchrotron X-ray diffraction. J Mol Biol 273, 729–739.

Tully BJ, Graham ED, and Heidelberg JF (2018). The reconstruction of 2,631 draft metagenome-assembled genomes from the global oceans. Scientific Data 5, 170203.

Uehling J, Deveau A, and Paoletti M (2017). Do fungi have an innate immune response? An NLR-based comparison to plant and animal immune systems. PLoS Pathog 13, e1006578.

Urbach JM, and Ausubel FM (2017). The NBS-LRR architectures of plant R-proteins and metazoan NLRs evolved in independent events. Proc Natl Acad Sci U S A 114, 1063–1068.

van der Biezen EA, and Jones JD (1998). The NB-ARC domain: a novel signalling motif shared by plant resistance gene products and regulators of cell death in animals. Current biology: CB 8, R226–227.

van der Meij A, Worsley SF, Hutchings MI, and van Wezel GP (2017). Chemical ecology of antibiotic production by actinomycetes. FEMS Microbiol Rev 41, 392–416.

Van Gerven N, Van der Verren SE, Reiter DM, and Remaut H (2018). The Role of Functional Amyloids in Bacterial Virulence. J Mol Biol 430, 3657–3684.

Wan W, and Stubbs G (2014). Fungal prion HET-s as a model for structural complexity and self-propagation in prions. Proc Natl Acad Sci U S A 111, 5201–5206.

Wang J, Hu M, Wang J, Qi J, Han Z, Wang G, Qi Y, Wang HW, Zhou JM, and Chai J (2019). Reconstitution and structure of a plant NLR resistosome conferring immunity. Science 364.

Wang SY, A JY, Fei F, Geng JL, Peng Y, Ouyang BC, Wang P, Jin XL, Zhao YQ, Wang JK, et al. (2017). Pharmacokinetics of the prototype and hydrolyzed carboxylic forms of ginkgolides A, B, and K administered as a ginkgo diterpene lactones meglumine injection in beagle dogs. Chin J Nat Med 15, 775–784.

Wang Y, and Jardetzky O (2002). Probability-based protein secondary structure identification using combined NMR chemical-shift data. Protein Sci 11, 852–861.

Wasmer C, Lange A, Van Melckebeke H, Siemer AB, Riek R, and Meier BH (2008). Amyloid fibrils of the HET-s(218-289) prion form a beta solenoid with a triangular hydrophobic core. Science 319, 1523–1526.

Wasmer C, Zimmer A, Sabate R, Soragni A, Saupe SJ, Ritter C, and Meier BH (2010). Structural similarity between the prion domain of HET-s and a homologue can explain amyloid cross-seeding in spite of limited sequence identity. J Mol Biol 402, 311–325.

Waterman MS, and Eggert M (1987). A new algorithm for best subsequence alignments with application to tRNA-rRNA comparisons. Journal of Molecular Biology 197, 723–728.

Wickner RB, Edskes HK, Gorkovskiy A, Bezsonov EE, and Stroobant EE (2016). Yeast and Fungal Prions: Amyloid-Handling Systems, Amyloid Structure, and Prion Biology. Adv Genet 93, 191–236.

Wozniak PP, and Kotulska M (2015). AmyLoad: website dedicated to amyloidogenic protein fragments. Bioinformatics 31, 3395–3397.

Xue JY, Wang Y, Wu P, Wang Q, Yang LT, Pan XH, Wang B, and Chen JQ (2012). A primary survey on bryophyte species reveals two novel classes of nucleotide-binding site (NBS) genes. PLoS One 7, e36700.

Yeager CM, Gallegos-Graves LV, Dunbar J, Hesse CN, Daligault H, and Kuske CR (2017). Polysaccharide Degradation Capability of Actinomycetales Soil Isolates from a Semiarid Grassland of the Colorado Plateau. Applied and Environmental Microbiology 83, e03020–03016.

